# Robust Prediction of Enzyme Variant Kinetics with RealKcat

**DOI:** 10.1101/2025.02.10.637555

**Authors:** Karuna Anna Sajeevan, Abraham Osinuga, B Arunraj, Sakib Ferdous, Nabia Shahreen, Mohammed Sakib Noor, Shashank Koneru, Laura Mariana Santos-Correa, Rahil Salehi, Niaz Bahar Chowdhury, Randy Aryee, Brisa Calderon-Lopez, Ankur Mali, Rajib Saha, Ratul Chowdhury

**Affiliations:** Department of Chemical and Biological Engineering, Iowa State University, Ames, Iowa, USA; The Center for Biorenewable Chemicals, Iowa State University, Ames, Iowa, USA; Department of Chemical and Biomolecular Engineering, University of Nebraska-Lincoln, Lincoln, Nebraska, USA; Department of Computer Science and Engineering, University of South Florida, Tampa, Florida, USA

**Keywords:** Enzyme kinetics, Bio-aware Machine learning, Enzyme engineering, Biocatalysis, Database curation

## Abstract

Predicting enzyme kinetics directly from sequence remains a central challenge in computational biology, particularly in resolving the effects of mutations at catalytically essential residues. Existing models frequently overlook the functional consequences of such perturbations, often defaulting to wild-type predictions even in cases of substantial activity loss, thereby limiting their reliability for enzyme design and mechanistic inference. Here, we introduce RealKcat, a machine learning framework trained on KinHub-27k, a rigorously curated dataset of 27,176 experimentally reported enzyme–substrate entries consolidated from BRENDA, SABIO-RK, and UniProt and verified across 2,158 primary sources. To ensure biochemical realism, kinetic parameters were collapsed into order-of-magnitude bins, enabling predictions that are tolerant to experimental noise yet sensitive to functional shifts. RealKcat integrates ESM embeddings for enzyme sequences with ChemBERTa embeddings of affiliated substrate, producing a unified feature space of the chemical conversion that supports robust multi-class classification of both catalytic turnover (*k_Cat_*) and substrate affinity (*K_M_*). Across cross-validation, hold-out, out-of-distribution, and few-shot evaluations—including a dense mutational landscape of alkaline phosphatase (PafA)—RealKcat consistently capturead the direction and magnitude of mutation-induced changes, while preserving discrimination in both wild-type and mutant contexts. Importantly, structural descriptors were deliberately excluded, as naive integration of structural features has been shown to impair model generalization, underscoring the primacy of rigorous dataset curation, biologically informed task formulation, and balanced evaluation metrics. RealKcat establishes a scalable and mutation-sensitive framework for enzyme kinetics prediction, offering a biologically grounded platform for enzyme engineering, metabolic modeling, and therapeutic design.

**Significance Statement:** Enzymes catalyze biochemical reactions that sustain life, and accurate measurement of their efficiency—expressed through turnover number (*k_Cat_*) and substrate affinity (*K_M_*)—is fundamental to biotechnology, synthetic biology, and even pharmaceutical innovation. Yet experimental assays remain prohibitive, time-intensive, and sensitive to conditions such as pH, temperature, and ionic strength of the assay buffer, while existing computational approaches often lack sensitivity to catalytic-site mutations and are constrained by inconsistencies in public databases. RealKcat addresses these gaps by introducing a rigorously curated dataset (KinHub-27k) derived from manual review of 2,158 articles and augmented with 5,278 synthetic catalytic variants generated through alanine substitution at annotated catalytic residues. Leveraging protein and substrate embeddings and a classification scheme based on order-of-magnitude kinetic bins, RealKcat achieves state-of-the-art functional e-accuracy and, critically, demonstrates sensitivity to catalytic perturbations. By adopting e-accuracy—a performance metric that evaluates predictions within ±1 order of magnitude, aligning with the practical utility of enzyme kinetics—RealKcat provides biologically meaningful assessments that conventional metrics often obscure. This work establishes a robust, mutation-aware predictive platform that advances computational enzyme design and extends applicability to biomanufacturing, metabolic engineering, and precision medicine.

## Introduction

Precise characterization of enzyme kinetics is foundational for synthetic biology, systems biology, and disease biomarker discovery, where tailored enzymes drive innovations in biomanufacturing and therapeutics (*1*, *2*). Predicting catalytic activity enables rapid mapping of metabolic landscapes in non-model species, engineering of enzymes for specific substrates, and optimization of biochemical pathways, thereby reducing reliance on costly experimental screening. Such predictive capability is particularly valuable for pathway optimization—whether for sustainable metabolite production or for dissecting disease-linked metabolic shifts—where trial-and-error experimentation is prohibitive and does not reveal mechanistic underpinnings. Because functionally diverse enzymes are frequently linked to sequence variation, including engineered mutations and naturally occurring polymorphisms, robust and mutation-sensitive models are essential both for industrial biocatalysis and for identifying disease prognosis markers (*3*, *4*).

Current computational approaches, however, often lack the sensitivity to detect functionally relevant alterations at catalytically crucial residues (*5*). Compounding this challenge, most reported kinetic measurements are generated under *in vitro* conditions that do not capture the dynamic biochemical environments of cells, leading to predictions that can diverge from *in vivo* behavior. This gap limits the accurate modeling of metabolic shifts, drug responses, and synthetic pathway performance, particularly in non-model organisms and complex human systems. With growing demand for precision enzyme design in biotechnology, metabolic engineering, and biomedicine, there is a pressing need for machine learning (ML) frameworks that offer mutation-aware, catalytically sensitive predictions built on rigorously curated datasets.

Recent advances in ML have produced promising enzyme kinetics predictors, including DLKcat, TurNuP, UniKP, CatPred, and EITLEM-Kinetics, each demonstrating the utility of data-driven approaches in modeling enzyme–substrate kinetics (*6–10*). DLKcat applies convolutional and graph neural networks to predict turnover numbers (*k_Cat_*) across diverse pairs, though its performance depends heavily on dataset diversity. TurNuP extends this by employing ESM-1b sequence encodings (*7*, *11*) with RDKit-derived reaction fingerprints in a gradient-boosted algorithm, improving generalizability for poorly represented enzymes. UniKP incorporates environmental descriptors such as pH and temperature into a two-layer model that encodes sequences and substrates (*8*), though its accuracy is constrained by training data quality. CatPred leverages neural networks trained on SABIO-RK and BRENDA (*12*, *13*), achieving state-of-the-art accuracy with approximately 76% of *k_Cat_* predictions and 84% of *K_M_* predictions within one order of magnitude (*10*). EITLEM-Kinetics advances further by employing an ensemble iterative transfer learning approach to predict kinetic parameters for mutants with low sequence similarity (*14*). These models employ random splitting schemes to generate training, validation, and test sets; TurNuP additionally excluded overlapping enzyme sequences across splits, while CatPred enforced the stricter condition that no enzyme–substrate pair was duplicated across partitions.

While these tools mark significant progress, they exhibit critical limitations in mutation sensitivity. In particular, alanine substitutions at catalytic residues yield near-identical predictions across models, reflecting reliance on global sequence similarity rather than explicit catalytic awareness. Moreover, CatPred’s concatenation of substrate and cofactor SMILES into a composite reaction fingerprint overlooks the independent effects of distinct ligands, constraining interpretability. For enzyme engineering, mutation sensitivity and dataset quality are paramount: accurate prediction of catalytic consequences requires not only sophisticated ML frameworks but also carefully curated training resources with consistent treatment of both wild-type and mutant variants. To address these challenges, we present RealKcat, a framework that integrates rigorous curation with optimized gradient-boosted classifiers to deliver robust, mutation-sensitive predictions of enzyme kinetics. RealKcat is trained on the KinHub-27k dataset, comprising 27,176 experimentally validated entries from BRENDA and SABIO-RK and augmented with 5,278 synthetic inactive sequences generated by alanine substitution at UniProt-annotated catalytic residues (*15*, *16*). Dataset inconsistencies (n = 1,804) were resolved through direct review of 2,158 primary sources, ensuring fidelity of kinetic parameters, sequences, and substrate assignments. By combining ESM sequence embeddings, which capture evolutionary and biochemical context (*17*), with ChemBERTa substrate embeddings derived from isomeric SMILES (*18*), RealKcat constructs a rich feature representation that enhances functionally relevant predictive accuracy and establishes a robust foundation for catalytically sensitive enzyme modeling.

Unlike previous models, RealKcat frames enzyme kinetics’ prediction as a classification problem, clustering *k_Cat_* and *K_M_* values by orders of magnitude with dedicated clusters for extreme values, a strategy that captures functional relevance across diverse enzyme classes. This is particularly important because industrial-scale enzyme engineering processes largely concern about a range of order of magnitude (say, 10^5^ < *k_cat_* < 10^7^) of a specific enzyme variant rather than the exact numerical value. In addition, predicting a range of kinetic values enables their usage in constructing feasible metabolic models (as shown before (*19*)) allowing for differences *in vitro* to *in vivo* kinetic rates, to capture experimental phenotypes with fidelity (*20–22*).

RealKcat. is a mutation-aware, classification-based framework for enzyme kinetics prediction that integrates rigorous data curation, order-of-magnitude clustering, and mechanistically guided negatives with advanced protein and substrate embeddings. We demonstrate that RealKcat achieves robust generalization across diverse evaluation regimes, including wild-type and mutant test sets, out-of-distribution catalytic variants, and a high-throughput mutational landscape of alkaline phosphatase (PafA). By emphasizing e-accuracy as a biologically grounded metric alongside balanced statistical measures such as Matthews Correlation Coefficient (MCC), RealKcat delivers predictions that are both functionally interpretable and statistically conservative. Beyond benchmarking, we demonstrate that RealKcat is capable of capturing mutation-induced kinetic shifts at catalytically relevant residues, highlighting its potential utility for enzyme design, metabolic engineering, and biomedical applications where mutation sensitivity is essential.

## Results

### Building a Curated Benchmark for Enzyme Kinetics

To establish a high-quality foundation for RealKcat, we curated a dataset of 27,176 experimentally verified enzyme kinetics entries, meticulously sourced from SABIO-RK, BRENDA, and UniProt (see Fig. 1A). Each entry underwent a rigorous corroboration process, wherein we cross-checked 2,158 source articles to resolve 1804 inconsistencies in catalytic parameters (*k_Cat_*, *K_M_*), enzyme sequences, and substrate identity (see Table 2). For enzymatic reactions involving multiple substrates, we included separate entries for each substrate, effectively allowing the same enzyme to appear more than once in the dataset. This design acknowledges the standard assumption underlying many reported *in vitro* characterizations—that assays are performed under saturating concentrations of co-substrates, such that the measured kinetic parameter reflects dependence on a single varied substrate (*23*). By separating substrates explicitly, the model can learn substrate-specific contributions to catalytic parameters without conflating effects from experimental co-substrate conditions. The curated KinHub-27k dataset and accompanying code are available for download and inference (https://chowdhurylab.github.io/downloads.html).

**Fig. 1.**
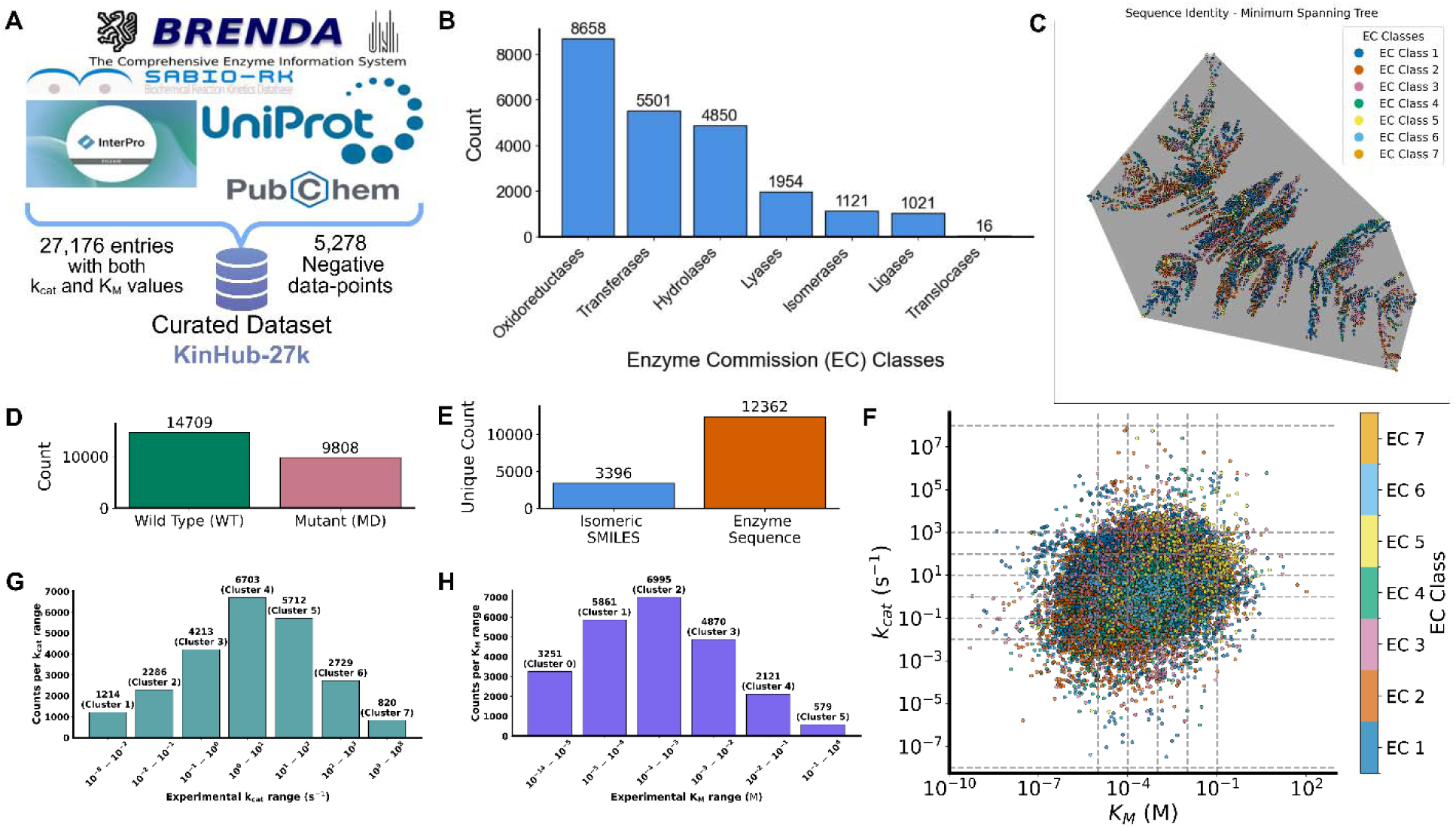
Comprehensive dataset curation, classification, and distribution for RealKcat model training. **(A)** Raw data for KinHub-27k was collated from multiple databases, including BRENDA, SABIO-RK, UniProt, and PubChem, resulting in a curated dataset with 27,176 entries containing both and *K_M_* values, and an additional 5278 synthetic inactive sequences **(B)** Distribution of enzyme entries across Enzyme Commission (EC) classes, highlighting a broad representation of enzymatic functions. **(C)** Network graph based on pairwise sequence identity, illustrating clustering by EC class and the diversity of sequence space represented in the dataset. **(D)** Counts of wild-type (WT) versus mutant (MD) entries, reflecting the dataset’s inclusion of both naturally occurring and engineered enzyme variants. **(E)** Unique counts of isomeric SMILES for substrates and enzyme sequences, indicating the dataset’s structural diversity. **(F)** Log-log scatter plot of experimental *K_M_* and values, depicting the core dynamic range of kinetic parameters. **(G & H)** Distribution of kinetic parameter clusters for and *K_M_*, respectively, categorized by order of magnitude to facilitate robust multiclass classification.

This intensive manual curation, addressing discrepancies that could impair the dataset’ reliability, enhances prediction accuracy for machine learning applications, making RealKcat the first enzyme kinetic predictor trained on a rigorously curated dataset. The curated dataset contains 3,396 unique SMILES, 12,362 sequences, and spans 1,499 unique organisms (Fig. 1E). Sequences longer than 1024 characters were removed, and duplicate entries were consolidated by grouping identical sequence and isomeric SMILES pairs. Rather than retaining the extreme (max–min) kinetic values—a common practice in prior models that can over-emphasize atypical or poorly replicated measurements—we observed in our own preliminary processing that this approach could produce spurious variability, with several unique sequence–substrate pairs differing by as much as 6–8 orders of magnitude (Supplemental Fig. S1). To mitigate this, duplicates were collapsed by taking the median *k_Cat_* and median *K_M_*, yielding robust central estimates that preserve the underlying biological signal while reducing the influence of outliers. This refinement produced a dataset of 24,517 unique entries with 14,709 wild-type entries and about 9808 mutant entries (Fig. 1D). Figure 1B shows the distribution of enzyme commission (EC) numbers, while Fig. 1C visualizes sequence identity–based relationships across enzymes. The network was constructed from pairwise global alignments, with distances defined as 1−identity, and simplified using a minimum spanning tree. Clustering by EC class highlights both conserved families and the broader diversity of sequence space encompassed in KinHub-27k. In order to strengthen the exposure of RealKcat functionally impaired sequences, we added 5278 mechanistically guided synthetic negative set by substituting UniProt-annotated “active-site” residues with alanine. In addition, to ensure robust target representations, duplicate sequence–substrate pairs were collapsed by median pooling prior to partitioning, a refinement that set the stage for reformulating kinetic prediction as a classification problem. Having established a curated dataset, we next reformulated kinetic prediction as a classification problem to better reflect biological interpretation.

### Log-Scaled Binning Towards Functional Catalytic Regimes

To address the intrinsic exponential variability of enzyme kinetic parameters, *k_Cat_* and *K_M_* were reformulated from continuous regression targets into biologically meaningful, log_10_-spaced bins, transforming the task into one of discriminating parameter ranges rather than predicting exact point estimates (Fig. 1F–H), with additional clusters for extreme values (see Materials and Methods for details). This design reflects how kinetic data are interpreted in practice—where order-of-magnitude resolution is often sufficient for functional inference—and imposes boundaries that separate distinct catalytic and binding regimes while reserving additional clusters for extreme values. By aligning prediction labels with magnitude intervals, the model is trained to focus on functionally relevant scale differences, thereby dampening the influence of experimental noise and assay-specific variability that persist even after median pooling of duplicate sequence–substrate pairs.

Moreover, the resulting formulation also enables the use of e-accuracy, a biologically grounded metric that rewards predictions within one bin of the measured value, acknowledging that slight deviations in magnitude rarely alter biochemical interpretation (*14*, *24*). This is particularly critical given that, even after rigorous curation, many enzyme–substrate pairs exhibit spreads of 6–8 orders of magnitude (Supplemental Fig. S1), driven by differences in organism source, assay protocols, and measurement conditions. In this framework, binning and e-accuracy operate synergistically to produce a stable, interpretable, and functionally relevant classification setting, strengthening RealKcat’s robustness for enzyme design and metabolic modeling applications.

### Synthetic Negatives Anchor Catalytic Loss Detection

To strengthen RealKcat’s ability to distinguish catalytically relevant changes, we expanded the dataset with a mechanistically guided synthetic negative set. This was generated by systematically substituting UniProt-annotated catalytic “active-sites” residues with alanine (see *Materials and Methods*), a perturbation most often expected to impair catalytic activity. This procedure yielded 5,278 high-confidence synthetic negatives, anchored by 22 experimentally curated WT–MD_Ala_ pairs that consistently exhibited substantial catalytic impairment, (Supplemental Table 1)—typically *≥*2 orders-of-magnitude reductions in *k_Cat_* and ≥1 order increases in *K_M_* for active site substitutions—without recourse to arbitrary zero-assignment policies. These curated examples provided benchmarks for calibrating synthetic negatives against realistic biochemical effects.

To further refine the set, we trained a binary meta-label classifier to discriminate between alanine substitutions that caused substantial catalytic loss and those that retained measurable activity. These curated WT–MD_Ala_ pairs served as the foundation for training on a mechanistically descriptor set, which included mutational severity (aggregate BLOSUM62 costs for alanine replacement), substrate physicochemical properties (molecular weight, logP, ring count, hydrogen-bond donor/acceptor counts), interaction-scale metrics (number of known active-site and substrate-to-enzyme molecular weight ratio), and the wild-type kinetic bin as a contextual reference. Under leave-one-out cross-validation, the classifier achieved accuracies of 77.27% for *k_Cat_* drop prediction (confusion: [5, 3; 2, 12]) and 81.82% for *K_M_* increase prediction (confusion: [4, 2; 2, 14]), providing sufficient discrimination policy to guide relabeling of synthetic negatives. Applied across the alanine-scanned dataset, this strategy produced 5278 high-confidence negatives. Importantly, these negatives remained sequence-close to their wild-type counterparts but were reassigned to kinetically divergent bins, creating hard counter examples that increased the discriminative challenge during model training. By integrating these negatives, we impose on RealKcat to learn sensitivity to catalytic relevance, with expected robustness in separating functional from non-functional variants during downstream evaluation.

### Minimizing Data Leakage Through Structured Partitioning

To provide an independent evaluation set beyond the cross-validation training and evaluation, we first established a randomized catalytic inference hold-out comprising 100 positive entries proportionally sampled from the wild-type (WT) and mutant-derived (MD) pools under a fixed random seed, thereby preserving the endogenous WT:MD ratio while ensuring independence from the training partitions. In parallel, 10 synthetic negatives were withheld from the negative test pool; these were generated by alanine substitution at annotated catalytic residues. Although their cognate wild-type sequences were allowed in the positive training partitions downstream, the corresponding negative sequences were never exposed during training, positioning them as in-distribution with respect to enzyme identity yet unseen as explicit negatives. This two-tier design enabled evaluation of catalytic inference across both biological variants and synthetic chemotypes, complementing the cross-validation partitions.

Beyond this hold-out, a careful manifold-guided clustering of the negative set yielded Affinity Propagation clusters, including a small “miscellaneous” bin, ensuring broad structural coverage while preventing over-representation of densely sampled regions (see Materials and Methods). This resulted in curated positives comprised 14,161 wild-type and 9,372 mutant entries after the randomized hold-out (based on median-pooled targets), forming the basis for subsequent modeling. Synthetic negatives were consistently paired with their cognate wild-type enzymes, and a 22-sample “anchor” synthetic set was withheld for drop-policy labeling of unlabeled synthetic negatives and for benchmarking. To preserve out-of-distribution integrity, sequence similarity between the anchor set and all other subsets (positives and negatives) was limited to ≤60% global identity. Across five folds, composite-label stratification preserved balanced kinetic class and WT:MD proportions, ensuring consistent distributions. Final train/validation/test proportions were tightly controlled (∼80/10/10 for CV folds and ∼95/2.5/2.5 for the negative set), supporting robust, leakage-minimized, and reproducible model evaluation.

### Sequence–Substrate Embeddings Capture Biochemical Context

With the inputs and target labels defined, we next compared two enzyme representation schemes. Specifically, we evaluated enzyme representations derived from two recent protein language models: ESM-2 and ESM-C. Both models, trained on large-scale sequence corpora, generate residue-level contextualized vectors that capture evolutionary and biochemical constraints across protein families (*11*, *17*, *25*). In this study, residue embeddings were aggregated into fixed-length vectors to serve as enzyme descriptors, a step that improves computational tractability for large-scale. For substrates, we used ChemBERTa embeddings (768 dimensions), derived from isomeric SMILES strings, which capture information related to stereochemistry, functional groups, and other physicochemical properties encoded in molecular structure (*18*). Enzyme and substrate embeddings were concatenated to form a 1,920-dimensional joint representation (1,920 dimensions for ESM-C as in Fig. 2A, 2,048 for ESM-2), integrating sequence context and substrate chemistry within a unified feature space. Although such concatenation does not explicitly model enzyme–substrate interactions, it provides a joint input space from which the classifier can infer associations between catalytic variation and molecular context. In addition, across random test and hold-out splits, ESM-2 and ESM-C performed comparably (e-accuracy, AUC-PR), whereas the main separation appeared in MCC on the out-of-distribution “anchor” set (n=22) and on a high-throughput alkaline phosphatase dataset (PafA) few-shot test, where ESM-C was higher (Fig. 3A). Given our emphasis on generalization under distribution shift and few-shot mutation testing, we report ESM-C as the default representation, while noting that ESM-2 yielded comparable performance across most splits.

**Fig. 2.**
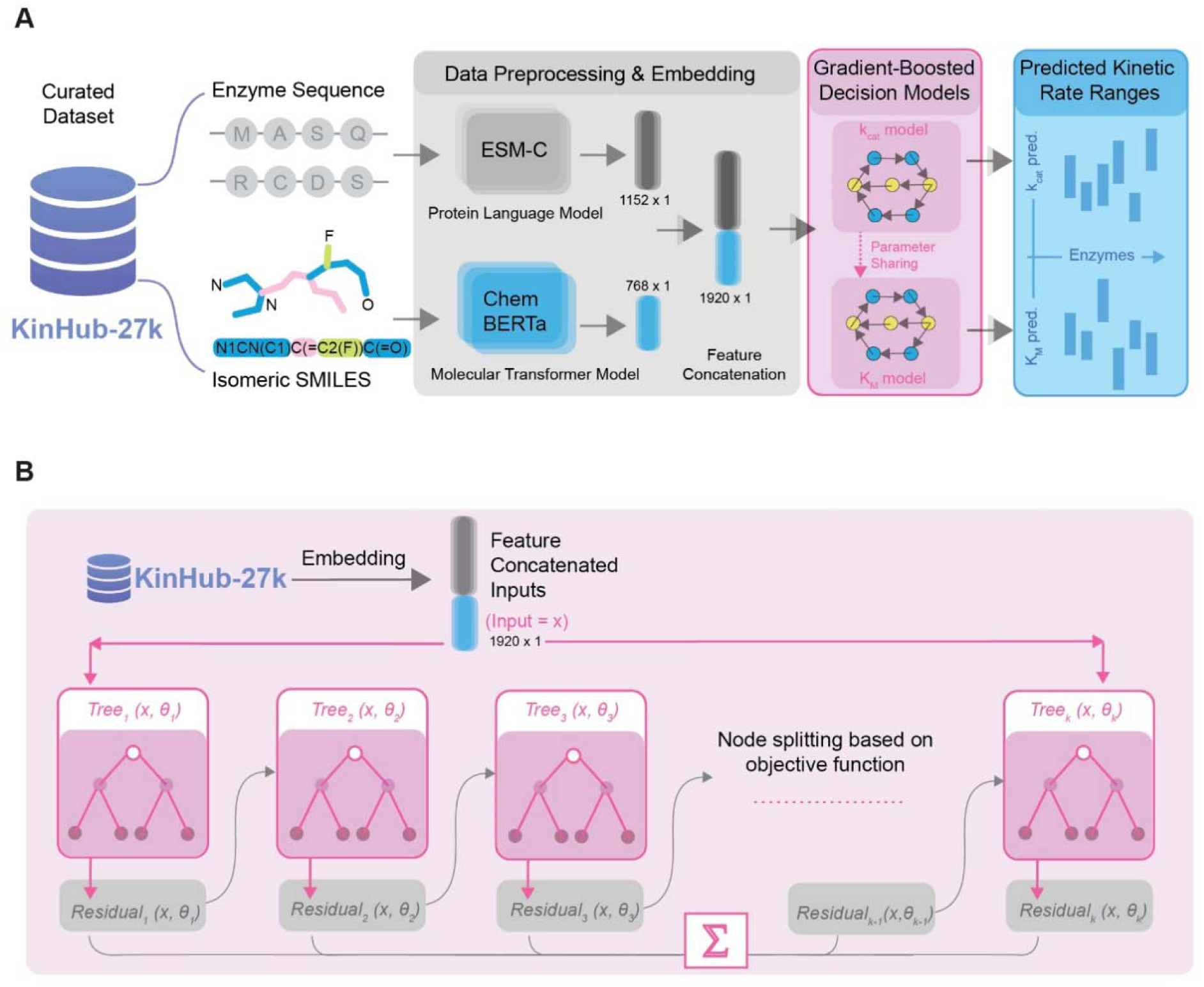
RealKcat model workflow and architecture for enzyme kinetics prediction. **(A)** The KinHub-27k dataset provides enzyme sequence and substrate information, embedded with ESM-C (1152-dimensional) and ChemBERTa (768-dimensional) models, respectively. The combined 1920-dimensional feature vectors are used to predict *k_Cat_* and *K_M_* values through separate gradient-boosted models, which share hyperparameters. Predictions are output as ranges, enhancing model stability across diverse enzyme-substrate pairs. **(B)** Gradient-boosted decision tree model architecture. Feature inputs are iteratively refined through an additive process of decision trees, each learning from the previous residuals to improve classification accuracy for predict *k_Cat_* and *K_M_* ranges

**Fig. 3.**
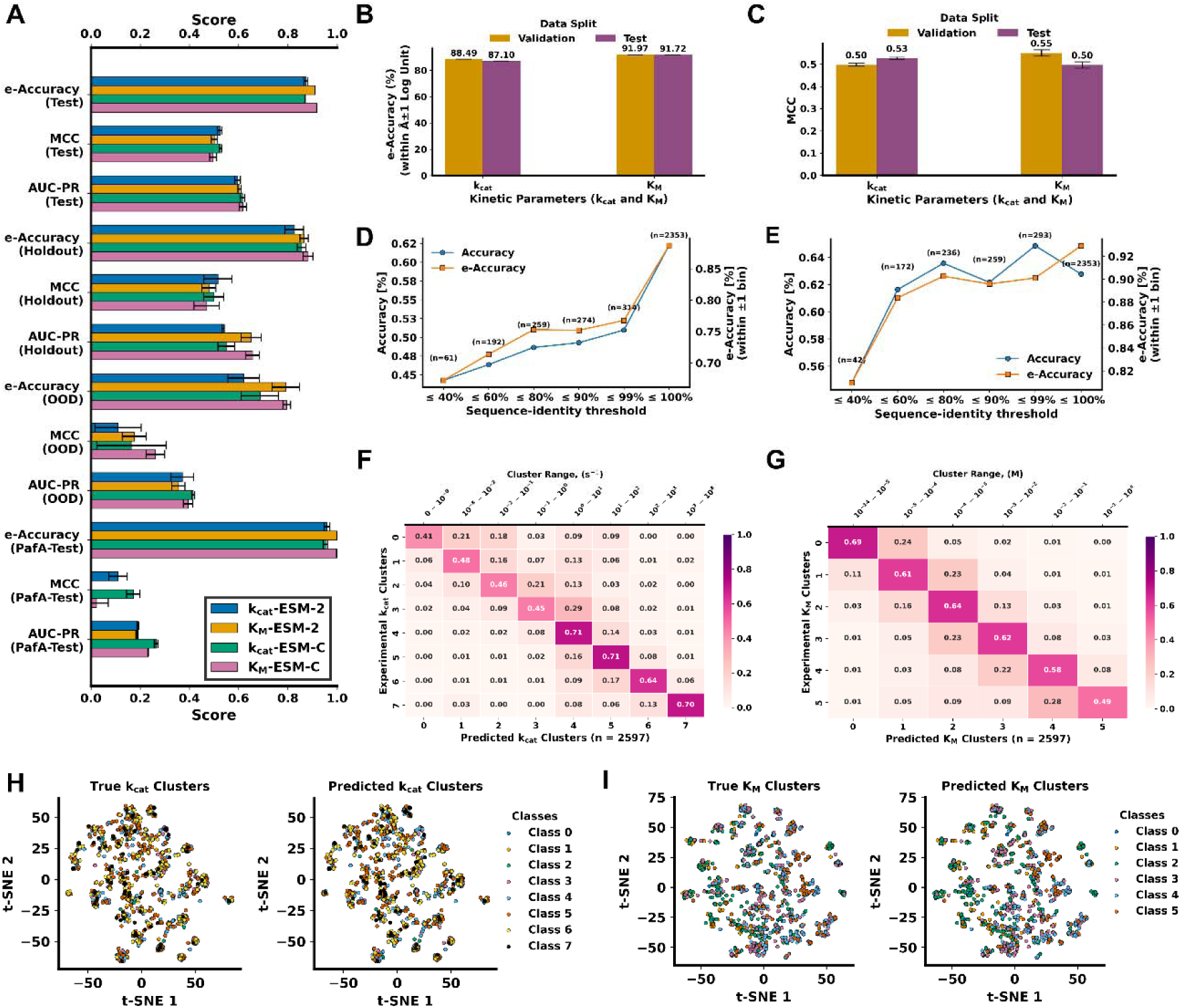
Evaluation of RealKcat performance across kinetic parameters and data splits. **(A)** Comparison of model performance using ESM-2 versus ESM-C embeddings for and *K_M_* across test, hold-out, out-of-distribution (OOD), and PafA few-shot splits, reported as e-accuracy, Matthews correlation coefficient (MCC), and area under the precision–recall curve (AUC-PR). **(B–C)** Performance consistency with ESM-C embeddings across validation and test sets: **(B)** e-accuracy (within ±1 log unit) and **(C)** MCC, demonstrating stable predictive capacity for both kinetic parameters. Panels A–C report the mean across all five folds ± standard deviation. **(D–E)** Generalization with ESM-C across sequence-identity thresholds for **(D)** and *K_M_* **(E)**. Accuracy and e-accuracy are plotted against sequence similarity to the training set, showing reliable extrapolation to low-identity regions. **(F–G)** Confusion matrices for **(F)** and **(G)** *K_M_* predictions, indicating strong recovery of experimental clusters with highest accuracy in central kinetic ranges and lower misclassification at extremes. **(H&I)** t-SNE visualizations of true vs. predicted clusters for and *K_M_*, showcasing high-dimensional separation and clustering.

### A Classification Framework for Enzyme Kinetics Prediction

RealKcat framework was trained using gradient-boosted additive modeling (XGBoost) to classify enzyme kinetic parameters into biologically meaningful log_10_-spaced bins (**Fig. 2B**). We emphasize from the outset that our objective was not to exhaustively search for an “optimal” algorithm but to demonstrate that, given a rigorously curated dataset and biologically grounded formulation, robust predictive performance can be achieved with a stable, interpretable, and reproducible classifier. Gradient boosting was chosen for its ability to model complex non-linear dependencies, its resilience to overfitting across heterogeneous feature spaces, and its transparent decision structure (*26*, *27*). This choice reflects a deliberate focus on robustness and reproducibility rather than marginal improvements in score maximization.

Model performance is summarized in **Fig. 3A–C**, which reports e-accuracy (within ±1 log unit) and macro-averaged MCC, a balanced classification metric that incorporates all true/false positives and negatives, across test, hold-out, out-of-distribution (OOD), and few-shot evaluations. On the test set, RealKcat achieved 87.10% e-accuracy for *k_Cat_* and 91.72% e-accuracy for *K_M_* (Fig. 3B), with corresponding MCC values of 0.53 for *k_Cat_* and 0.50 for *K_M_* (Fig. 3C). These metrics remained consistent across validation and hold-out partitions, underscoring that the model moderately generalizes beyond its training distribution. Meanwhile, to address the strong class imbalance across kinetic bins, we applied SMOTE oversampling (*28*) exclusively to the training partition of each cross-validation fold, while leaving validation and test sets untouched to ensure fair generalization. This balancing step prevented majority-class dominance, stabilized performance in minority clusters, and directly contributed to the consistently high macro-MCC observed in training (Fig. 3C). Moreover, *K_M_* predictions exhibit consistently higher e-accuracy than *k_Cat_* (91.72% vs 87.10% on test set), reflecting the fact that substrate affinities cluster within narrower log10-ranges (Fig. 1F), whereas turnover numbers span broader, assay-sensitive dynamic ranges (Supplemental Fig. S1). This highlights the inherent difficulty of predicting catalytic rates and underscores that RealKcat conforms to the established hierarchy of predictability, with *K_M_* values being inherently more tractable than *k_Cat_*.

While e-accuracy captures the biological and environmental variational reality that order-of-magnitude predictions suffice for most functional inferences, MCC provides a balanced statistical measure that accounts for all entries in the confusion matrix and prevents dominance by majority classes. This is especially critical given the natural skew of enzyme kinetics data toward central bins (Fig. 1G-H). Exact-bin accuracy, F1-scores, recall, and area under the precision–recall curve (AUC-PR, which is more informative than ROC curves under class imbalance), are reported in **Supplemental Fig. S2**, where lower values illustrate the limitations of conventional metrics under class imbalance. In contrast, the stability of e-accuracy and MCC across splits (Fig. 3A–C) highlights why these measures are more conservative and robust indicators of predictive quality.

Prediction under distribution shift was further assessed on an OOD anchor set of 22 negatives and a few-shot alkaline phosphatase (PafA) dataset. In these cases, MCC slightly distinguished ESM-C from ESM-2, with modest but consistent improvements for ESM-C (Fig. 3A). Although performance across other metrics was comparable, these results emphasize the relevance of embedding choice for prediction in sparse or low-data settings, where robustness rather than average accuracy is most critical. Generalization across homology boundaries also supports this interpretation: stratification of predictions by sequence identity (Fig. 3D–E) shows that e-accuracy remain fairly stable down to 60% identity relative to training, with only modest declines below 40%. This indicates that the embedding framework captures catalytic information beyond sequence similarity, enabling reliable extrapolation to enzymes not represented in the training set.

Resolution at the class level is evident from the confusion matrices for *k_Cat_* and *K_M_* (**Fig. 3F–G**). Predictions are concentrated along the diagonal, with strongest agreement in the central bins but crucially, even in sparsely populated extremes, most predictions fall within ±1 log unit of the experimental cluster. This validates the use of e-accuracy as a biologically relevant measure, as near-miss predictions at the order-of-magnitude level still preserve functional interpretation. The t-SNE projections (Fig. 3H–I) provide further confirmation, with predicted classes reproducing the global structure of experimental clusters in reduced space, demonstrating that RealKcat has internalized meaningful distinctions across kinetic regimes rather than collapsing toward majority classes. For deployment, we report results from fold-5, pre-specified as the inference model, while population-level reproducibility is summarized as cross-fold mean ± SD.

### Mutation Sensitivity Across Catalytic Residues

RealKcat demonstrated sensitivity to point mutations at catalytically relevant residues, capturing the expected shifts in kinetic parameters when active-site positions are perturbed. To evaluate mutation awareness under controlled but distinct conditions, we adopted a two-tier scheme. First, we assessed performance on the established out-of-distribution (OOD) anchor set comprising 22 experimentally characterized wild-type (WT) and alanine variants at UniProt-annotated catalytic residues that were withheld from training. Second, we evaluated a negative hold-out consisting of 10 synthetic alanine variants drawn at random from the partitioned negative test pool; their cognate WT enzymes appeared in the positive training partitions, but the exact negative sequences were never used for training. This design probes two distinct generalization regimes: (i) an OOD setting in which both enzyme–substrate pairs and mutation outcomes are unseen, and (ii) an in-distribution-by-identity setting in which the enzyme family is familiar but the negative examples are novel, enabling an appraisal of mutation sensitivity independent of trivial sequence memorization.

Across the 22 OOD WT–MD*_Ala_* pairs (Supplemental Table 1), RealKcat reproduced both the directionality and order-of-magnitude shifts in catalytic impairment for *k_Cat_* and *K_M_*, with the majority of mutant predictions falling within ±1 bin of the experimental values (Fig. 4A–B). These pairwise comparisons show that predicted WT→MD trajectories align with experimentally observed changes, particularly for direct-contact substitutions at annotated catalytic residues. Longitudinal visualizations (Fig. S3) further illustrate this behavior, with predicted trajectories closely tracking experimental curves and remaining largely within the ±1-bin tolerance bands for both parameters. A critical observation, however, is that misclassified mutants tend to be predicted near the corresponding wild-type bin, indicating a fallback toward baseline activity rather than spurious overestimation of catalytic loss. This pattern highlights both the conservative nature of the model and the difficulty of extrapolating large mutational effects in an out-of-distribution regime. Together, these analyses indicate that RealKcat captures mutation-induced movement across kinetic bins for catalytic residues in a setting that is intentionally distribution-shifted from training.

**Fig. 4.**
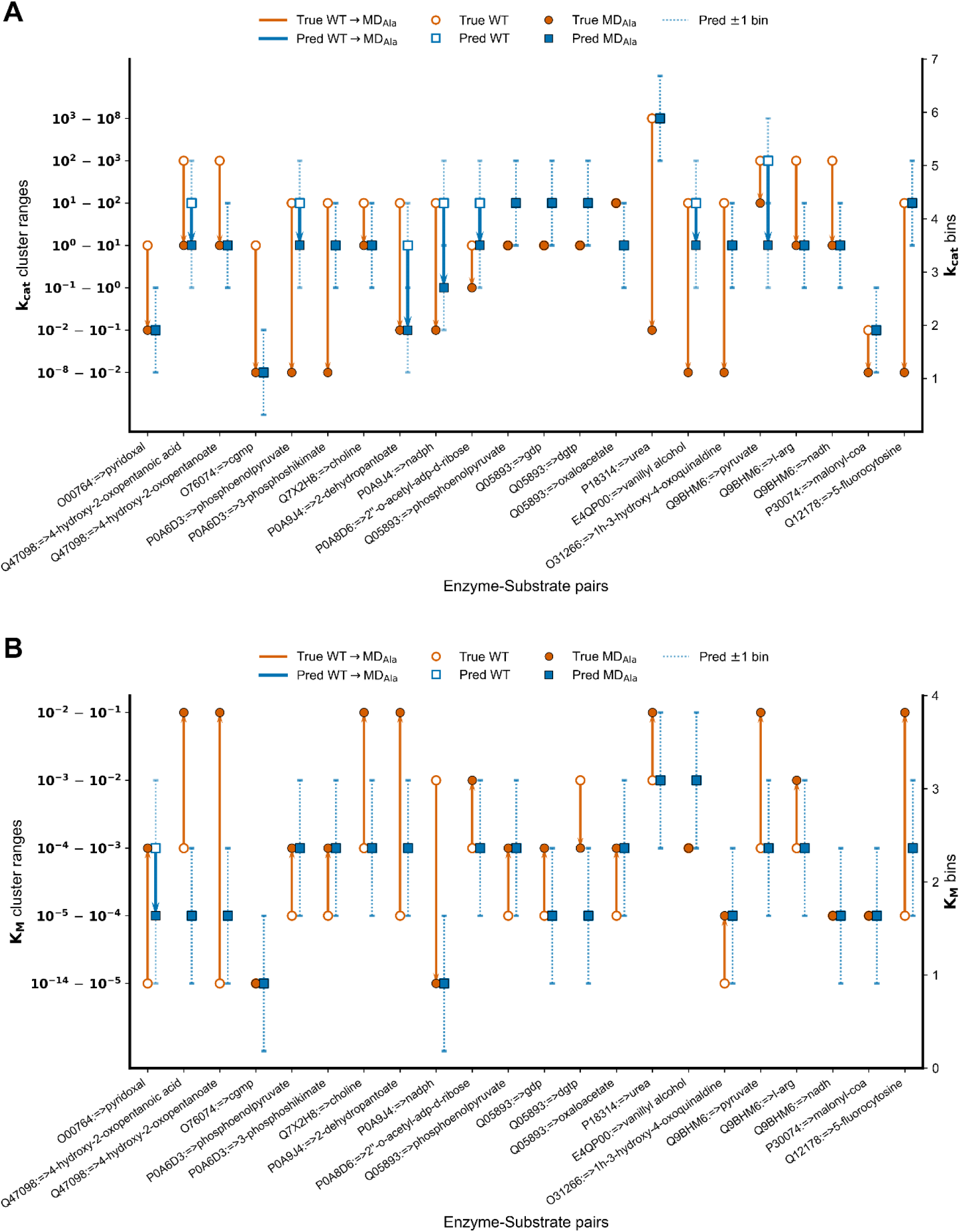
Out-of-distribution anchor evaluation on experimentally validated WT–MD*_Ala_* pairs at catalytic residues. **(A)** Predictions of clusters for 22 WT–MD*_Ala_* pairs curated from UniProt-annotated catalytic residues. Orange markers indicate experimentally measured wild-type (WT, open circle) and mutant (MD*_Ala_*, filled circle) values; blue markers denote RealKcat predictions for WT (open square) and MDAla (filled square). Lines connect WT–MD*_Ala_* pairs to illustrate mutation-induced shifts. Dotted vertical bars indicate the ±1 bin tolerance window used in e-accuracy assessment. Predicted shifts largely align with experimental measurements, with most mutant values falling within ±1 order of magnitude of the experimental cluster. **(B)** Equivalent analysis for *K_M_* clusters across the same 22 WT–MD*_Ala_* pairs. RealKcat predictions capture the direction and magnitude of affinity changes following alanine substitution at catalytic residues, with the majority of mutant values within ±1 bin of experimental measurements.

For the negative hold-out of 10 synthetic alanine variants (Table 1), which included not only single-point substitutions but also several dual-mutant cases (samples 2, 6, and 10), mutation effects were benchmarked against a binary meta-label (drop vs. no-drop for *k_Cat_*; increase vs. no-increase for *K_M_*) derived from mechanistic descriptors (*Materials and Methods*). On this set, RealKcat achieved precision = 1.00 and recall = 0.80 for *k_Cat_* drops (F1 = 0.87; MCC = 0.51) and precision = 1.00 and recall = 0.40 for *K_M_* increases (F1 = 0.57; MCC = 0.50), with corresponding confusion matrices shown in Fig. 5A–B. We further examined how well RealKcat could predict categorical *k_Cat_* and *K_M_* bins for alanine substitutions at all known catalytic residues, including those involving simultaneous perturbation at more than one active-site position. In scatter plots comparing predicted and experimental values, RealKcat’s wild-type predictions (Fig. 5C, E) aligned closely with experimental values and were largely within ±1 log bin of the identity line, comparable to other available zero-shot inference models (DLKcat, CatPred, UniKP, and EITLEM-Kinetics). However, we emphasize that RealKcat is not intended as an algorithmic competitor to prior deep-learning frameworks such as DLKcat or CatPred, but rather as a dataset-and evaluation-driven advance. While these published models are included as benchmarks to reflect mechanistic expectations, all primary performance claims for RealKcat are confined to standardized evaluation settings within KinHub-27k.

**Fig. 5.**
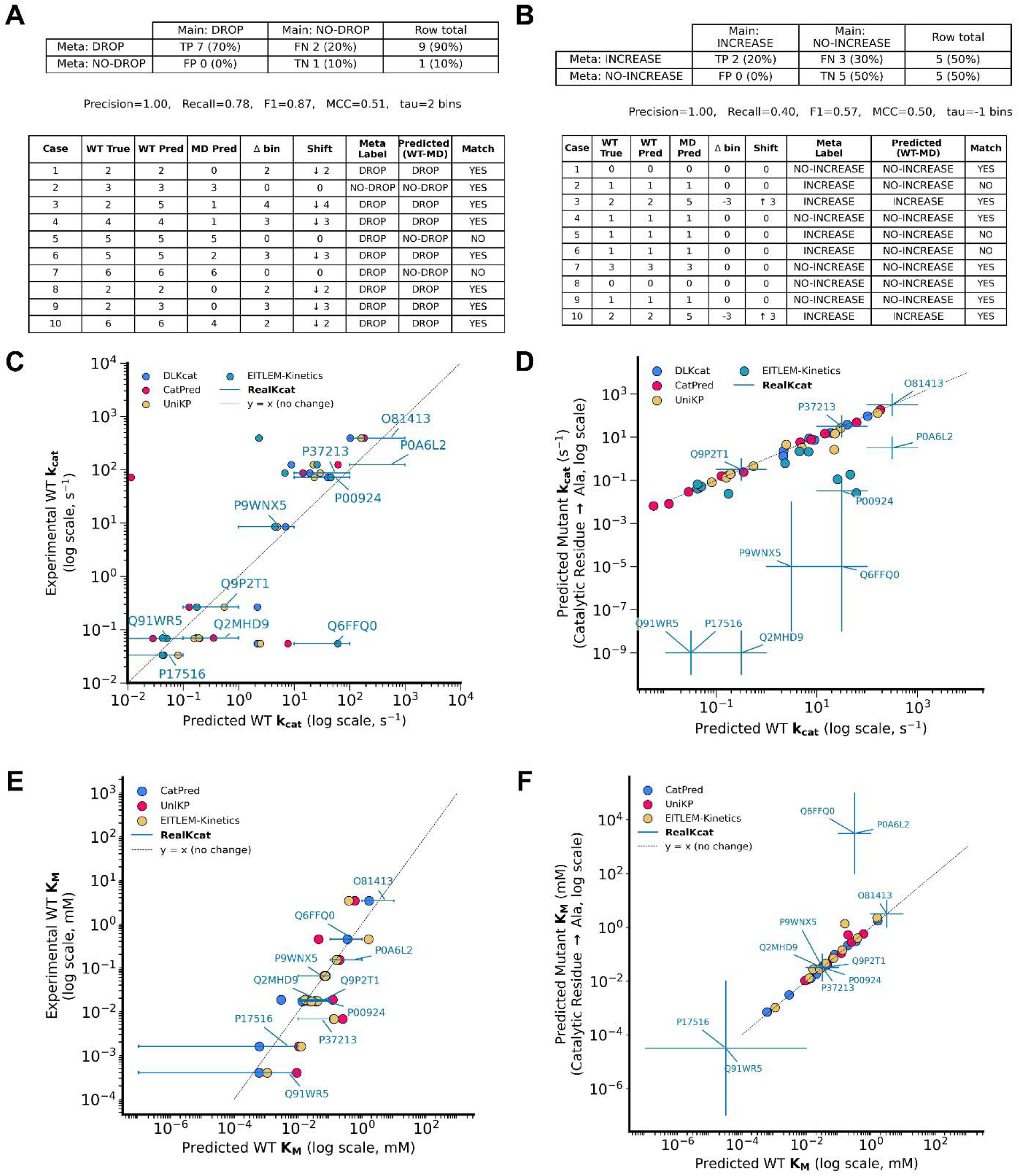
Evaluation of mutation-aware predictions on a synthetic negative hold-out of 10 catalytic-site alanine variants. **(A–B)** Confusion matrices and summarizing binary classification of catalytic loss versus retention across 10 held-out synthetic alanine variants. **(A)** “DROP” for *k_Cat_*, **(B)** “INCREASE” for *K_M_*. Predictions were benchmarked against meta-labels derived from mechanistic descriptors. **(C–F)** Scatter plots comparing predicted versus experimental kinetic values for wild-type (WT) and mutant (MD*_Ala_*) enzymes. **(C–D)** *k_Cat_* values for WT **(C)** and MD*_Ala_* mutants **(D)**. **(E–F)** *K_M_* values for WT **(E)** and MD*_Ala_* mutants **(F)**. The diagonal line denotes the identity relationship (x = y). In panels C–F, RealKcat predictions are shown as blue line segments connecting the corresponding experimental and predicted values. The length of each line segment denotes the span of the predicted log-bin cluster, while the direction of shift away from the identity line in panels D and F indicates whether the model predicted a decrease (leftward/downward) or increase (rightward/upward) relative to experimental values. These line segments are unique to RealKcat and should not be conflated with either the dotted identity line (x = y), which indicates perfect agreement, or the markers of other models (DLKcat, CatPred, UniKP, EITLEM-Kinetics), which are displayed as colored points without connecting segments.

**Table 1.**
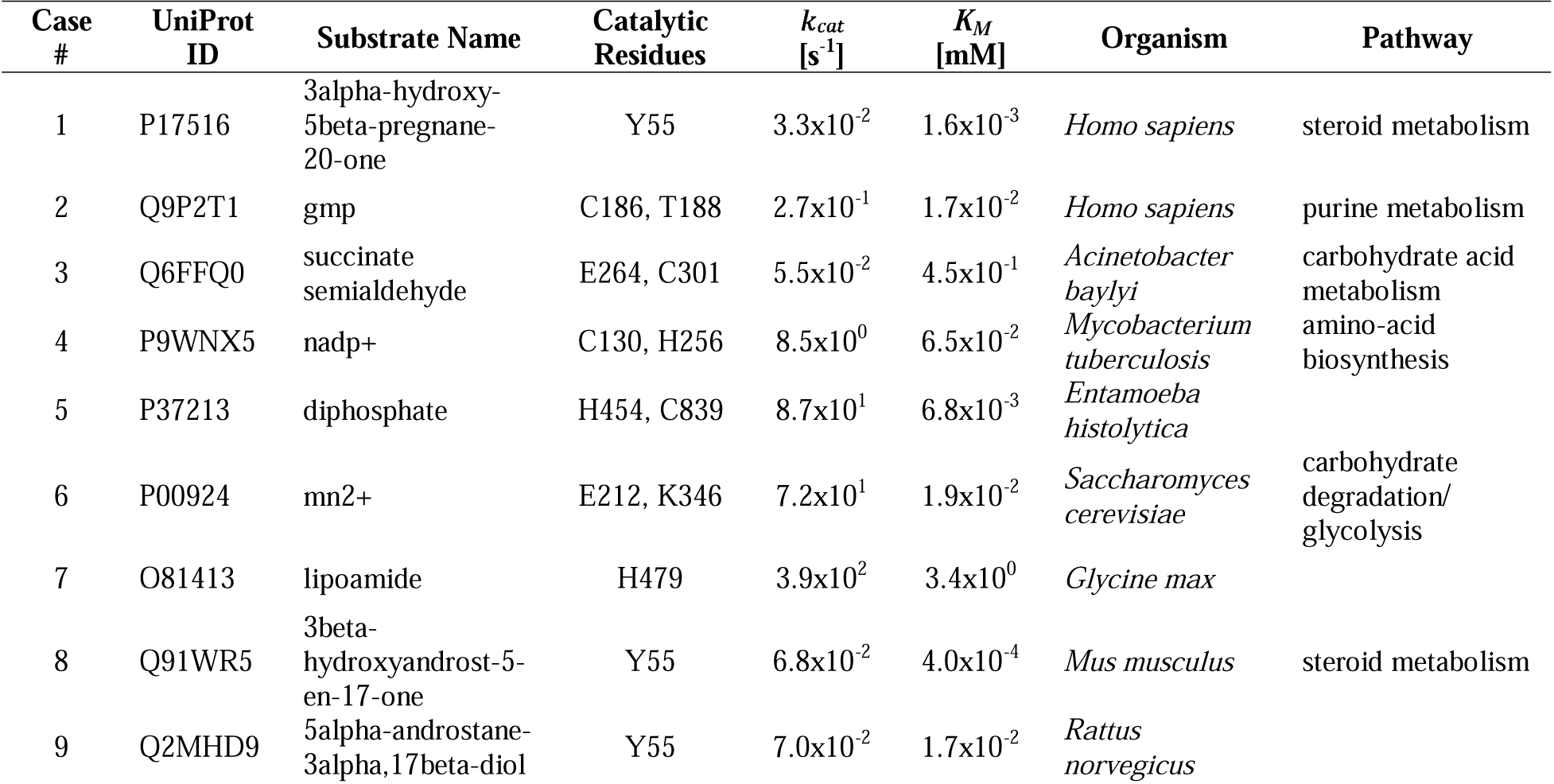
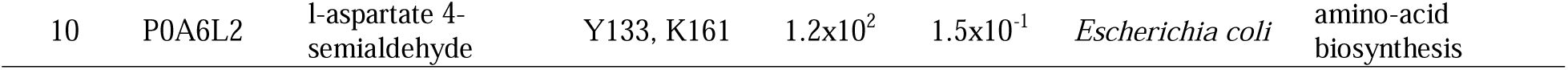
Independent in-distribution synthetic negative hold-out set of 10 catalytic-site alanine variants. Summary of wild-type (WT) enzymes whose corresponding alanine-substituted mutants used for in-distribution hold-out evaluation. These MD variants are excluded from training to provide a controlled evaluation of RealKcat’s mutation sensitivity in an in-distribution-by-identity regime.

| Case # | UniProt ID | Substrate Name | Catalytic Residues | $k_{cat}$ [s<sup>-1</sup>] | $K_M$ [mM] | Organism | Pathway |
| --- | --- | --- | --- | --- | --- | --- | --- |
| 1 | P17516 | 3alpha-hydroxy-5beta-pregnane-20-one | Y55 | $3.3 \times 10^{-2}$ | $1.6 \times 10^{-3}$ | *Homo sapiens* | steroid metabolism |
| 2 | Q9P2T1 | gmp | C186, T188 | $2.7 \times 10^{-1}$ | $1.7 \times 10^{-2}$ | *Homo sapiens* | purine metabolism |
| 3 | Q6FFQ0 | succinate semialdehyde | E264, C301 | $5.5 \times 10^{-2}$ | $4.5 \times 10^{-1}$ | *Acinetobacter baylyi* | carbohydrate acid metabolism |
| 4 | P9WNX5 | nadp+ | C130, H256 | $8.5 \times 10^0$ | $6.5 \times 10^{-2}$ | *Mycobacterium tuberculosis* | amino-acid biosynthesis |
| 5 | P37213 | diphosphate | H454, C839 | $8.7 \times 10^1$ | $6.8 \times 10^{-3}$ | *Entamoeba histolytica* | |
| 6 | P00924 | mn2+ | E212, K346 | $7.2 \times 10^1$ | $1.9 \times 10^{-2}$ | *Saccharomyces cerevisiae* | carbohydrate degradation/ glycolysis |
| 7 | O81413 | lipoamide | H479 | $3.9 \times 10^2$ | $3.4 \times 10^0$ | *Glycine max* | |
| 8 | Q91WR5 | 3beta-hydroxyandrost-5-en-17-one | Y55 | $6.8 \times 10^{-2}$ | $4.0 \times 10^{-4}$ | *Mus musculus* | steroid metabolism |
| 9 | Q2MHD9 | 5alpha-androstane-3alpha,17beta-diol | Y55 | $7.0 \times 10^{-2}$ | $1.7 \times 10^{-2}$ | *Rattus norvegicus* | |

**Table 2.** Resolution and additions in the KinHub-27k mutant dataset. This table details the curation process, highlighting resolutions and removals across parameters (*K_M_* and *k_Cat_*) substrates, and mutations. 1804 discrepancies were resolved, with 1,072 new entries added, refining the dataset to 11,175 mutant entries for predictive modeling.

| Parameter | corrected | removed |
| --- | --- | --- |
| $K_M$ | 788 | 10 |
| $k_{cat}$ | 618 | 10 |
| substrates | 240 | 8 |
| mutations | 18 | 21 |
| duplicates | - | 91 |
| additional data | 1072 | - |

When extended to catalytic-site mutants, RealKcat captured both the magnitude and direction of expected (meta-labeled) mutation-induced changes. For *k_Cat_*, MD*_Ala_* mutant predictions (Fig. 5D) consistently deviated from their wild-type cognates in accordance with the drop versus no-drop meta-labels, reproducing the separation pattern observed in the categorical evaluation (Fig. 5A). For *K_M_* most MD*_Ala_* mutant predictions (Fig. 5F) tracked the expected increase versus no-increase trends, again maintaining separation between perturbed and unaltered variants. In general, the scatter plots (Fig. 5C–F) extend the categorical observations of Fig. 5A–B into the continuous prediction domain, illustrating that the binary drop/increase labels are reflected in RealKcat’s predicted kinetic clusters. Although the evaluation set (n=10) is intentionally small to enforce strict hold-out conditions, these results demonstrate that RealKcat can distinguish major activity losses from retention even in an in-distribution-by-identity regime, where enzyme families are familiar but the explicit negative variants themselves remain unseen.

Taken together, the OOD anchor and in-distribution negative evaluations provide complementary perspectives: in both regimes RealKcat preserves the direction of WT→mutant effects and maintains bin-level agreement with experimental outcomes. Moreover, this behavior extends to non-alanine and broader OOD mutants (Fig. S4), further underscoring that model generalization is not limited to alanine substitutions. We stress that this section evaluates mutation sensitivity—that is, whether predicted movements across kinetic bins are consistent with perturbations at catalytic residues—rather than fine-grained ranking within bins. As expected for an order-of-magnitude classification framework, sub-bin resolution is not achieved; nevertheless, the observed separations provide convergent evidence that RealKcat internalizes biochemical constraints at catalytic sites and generalizes these constraints across distinct evaluation conditions.

### Few-Shot Generalization in a High-Throughput Mutational Landscape

To extend mutation-aware testing beyond curated catalytic residues and synthetic negatives, we employed the high-throughput dataset of the alkaline phosphatase (PafA) from *Elizabethkingia meningoseptica*, originally reported by Markin et al. (*29*). This dataset systematically introduces single-point mutations (glycine, valine and alanine) across the enzyme sequence to probe catalytic and structural roles, yielding kinetic measurements (*k_Cat_* and *K_M_*) for 1,016 variants with the substrate carboxy 4-methylumbelliferyl phosphate ester (cMUP), a binding-limited substrate. For evaluation with RealKcat, we partitioned the dataset into 398 training, 310 validation, and 310 test variants, ensuring that the training distribution spanned the high, medium, and low ranges of observed turnover values. Crucially, while the wild type (WT) and a catalytic-site variant (T79S) were present in the training set, the key catalytic mutations R164A and R164G—located at a residue central to substrate coordination (Fig. 6A)—were held out for the test split to specifically assess inferencing on unobserved but catalytically significant mutations.

**Fig. 6.**
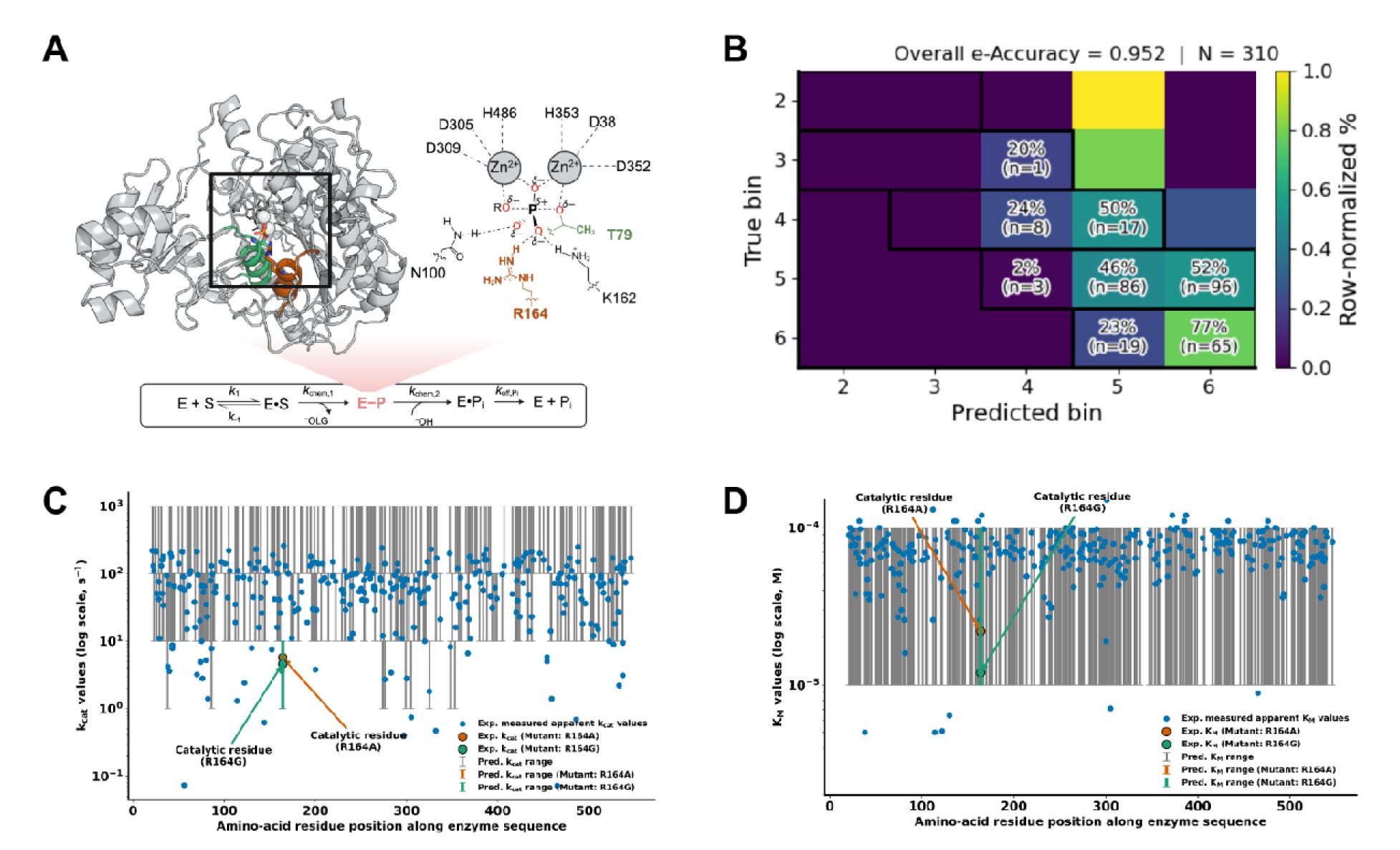
Few-Shot Learning Evaluation of RealKcat’s catalytic relevance prediction and sensitivity to mutations of alkaline phosphatase. **(A-D) (A)** Structure of alkaline phosphatase PafA with highlighted catalytic residues in transition state, showing positions T79 and R164 used in mutant testing. **(B)** Confusion-matrix-style visualization of predicted versus true bins (N = 310). Each cell shows the row-normalized percentage and the number of samples in parentheses; the overall ± 1 bin e-accuracy = 0.952. No single class dominates, confirming balanced coverage across bins 3–6. **(C-D)** Sequence-wide distribution of **(C)** and *K_M_* **(D)** values across the PafA test dataset, with RealKcat predictions compared to experimental data, specifically for key R164A and R164G mutations.

Across the PafA test set, RealKcat achieved 51% exact-bin accuracy for *k_Cat_* and 94.2% for *K_M_*, with markedly higher e-accuracy values of 95.2% (*k_Cat_*) and 100% (*K_M_*) within ±1 order of magnitude (Fig. 6C–D). As shown in Fig. 6B, the PafA test subset spans multiple kinetic bins with no single class dominating the distribution, confirming that these results are not attributable to class imbalance or majority-class effects. At the residue level, RealKcat correctly reproduced the bin-level placement and directional shifts associated with catalytic substitutions: both R164A and R164G were predicted within the same experimental bins as their *in vitro* measurements (Fig. 6C–D), with R164A and R164G showing marked reductions in turnover. This agreement underscores the model’s sensitivity to side-chain chemistry and conformational flexibility at catalytic sites. Nevertheless, we also observed occasional off-target predictions for other mutations, in which predicted values deviated from experimental bins without systematic bias and not within ±1 order of magnitude of experimental bins. Structural analysis suggests these discrepancies may arise from the dense network of contacts at these off-target residues, for instance H486 is highly interconnected with at least 17 neighboring residues (see Fig. S5). Perturbations at this residue are therefore likely to propagate non-locally, complicating predictive accuracy. At the same time, a recurring pattern was observed in cases of misclassification: when RealKcat failed to capture the mutation-induced impairment, predictions often fell back to a conservative baseline near the wild-type bin, typically within ±1 log unit of the WT value. This behavior reflects a conservative bias that prioritizes stability over speculative large deviations, but it also reveals where mutation effects may be underestimated.

To probe these failures systematically, we scanned for mutants experimentally reported to exhibit ≥2-bin drops from the WT reference bin (e.g., WT at bin 6, mutant ≤ bin 3) and then examined whether RealKcat could recover these severe impairments within ±1 log-bin tolerance. Strikingly, only D305 was predicted correctly within ±1 bin of the experimental cluster, while the majority of other severe-drop mutants were misclassified back toward the wild-type reference, reflecting a conservative fallback bias. These failures clustered primarily at loop-like or disordered regions, or at structurally dense positions where ≥10 residues fall within a 6 Å radius. Examples include G56, G465, L297, D144, F332, and H486—sites embedded in flexible or highly interconnected regions where mutation-induced perturbations likely propagate in ways not captured by sequence embeddings alone. In such contexts, RealKcat underestimates the magnitude of catalytic loss, highlighting its current limitation in modeling destabilizing mutations at disordered or densely packed sites without explicit structural context.

Importantly, this analysis underscores the value of ±1 log-bin e-accuracy as a physiologically relevant measure. By maintaining predictive fidelity in such few-shot settings, RealKcat demonstrates generalizability to densely sampled mutational landscapes, complementing its performance on curated OOD anchors and synthetic negatives. Moreover, experimental enzymatic turnover at catalytic residues can vary by up to an order of magnitude across assay conditions due to effects of pH, ionic strength, temperature, and protonation states (Fig. S1). Within this biological variability, RealKcat consistently reproduced the qualitative directionality and quantitative bin-level placement of PafA mutations.

## Discussion

RealKcat provides a mutation-sensitive, catalytically aware framework for predicting enzyme kinetics by integrating rigorous dataset curation, pretrained biochemical language models, and a robust classification-based modeling scheme. Rather than relying on methodological novelty at the algorithmic level, our approach demonstrates how careful attention to data quality, representation, and biologically grounded evaluation can substantially improve reliability and generalizability. The KinHub-27k dataset, built through meticulous reconciliation of thousands of source articles and manual resolution of inconsistencies, underpins this framework and distinguishes it from prior enzyme kinetics resources. Median pooling of duplicates, mechanistically guided generation of negatives, and stratified data partitioning together establish a dataset and modeling strategy that is resistant to spurious variability and leakage—issues that often obscure performance in regression-based formulations. By recasting *k_Cat_* and *K_M_* as order-of-magnitude classification tasks, RealKcat addresses the intrinsic variability of kinetic measurements and provides predictions in physiologically interpretable ranges, bridging the gap between *in vitro* assay heterogeneity and *in vivo* functional relevance (*24*).

Moreover, RealKcat’s dual-input formulation is not an architectural embellishment but a biological necessity. Kinetic parameters such as *k_Cat_* and *K_M_* arise from the coupled enzyme– substrate context; enzyme-only or substrate-only models cannot resolve this interdependence and therefore underperform when tasked with catalysis-specific predictions. By grounding our modeling unit in enzyme–substrate pairs, RealKcat aligns computational inference with the biochemical basis of catalysis, avoiding the pitfalls of single-input formulations more appropriate for binding screens than for turnover and affinity estimation. Importantly, most experimental kinetic assays are performed under saturating concentrations of co-substrates and cofactors, such that reported *k_Cat_* and *K_M_* values are conditional on the specific enzyme–substrate pair being assayed. This experimental design makes it biologically inconsistent to decouple enzyme from substrate in predictive modeling: a single-input model cannot capture the conditional dependency imposed by assay context. Hence, single-input ablations were considered out-of-scope for RealKcat, as they would conflict directly with our rationale and reduce biological interpretability.

Furthermore, the classification-based formulation adopted in RealKcat is not merely a technical simplification but a deliberate reflection of how kinetic data are interpreted in practice. Turnover numbers and affinities are rarely known with precision beyond orders of magnitude, and even for the same enzyme can vary by several bins, as much as three orders of magnitude, depending on experimental context (*1*, *24*, *30*). For instance, according to criteria from Bar-Even et al. (*24*), *K_M_* values can be categorized into high (>1 mM), moderate (0.1–1 mM), and low (≤0.1 mM) clusters, distinguishing enzymes that maintain metabolic flux across dynamic conditions from those requiring high substrate affinity. Similarly, *k_Cat_* values span high (>1000 s^−1^), moderate (100–1000 s^−1^), and low (≤100 s^−1^) categories, reflecting known variation in catalytic efficiency. Moreover, *in vivo* data themselves can be noisy and context-dependent due to the temporal variability and regulatory dynamics inherent to living systems (*31*, *32*), complicating their direct interpretation.

By training the model to operate at this resolution, we emphasize robustness and functional interpretability over pointwise accuracy. The use of e-accuracy as a central evaluation metric acknowledges this reality, rewarding predictions within ±1 order of magnitude of experimental values, while macro-averaged MCC ensures balanced evaluation across abundant and rare kinetic classes. Together, these measures prioritize stability and biological meaning over superficial improvements in regression metrics. This evaluation strategy also clarifies why apparent “failures” often reflect a conservative bias: when RealKcat misclassifies, predictions frequently fall back toward wild-type activity rather than producing implausible deviations, underscoring both its strengths and its current limitations. Technically, from the inputs, residue-level embeddings were aggregated via mean pooling to yield fixed-length descriptors that capture conserved evolutionary signals, sequence-context dependencies, and biochemical constraints across enzyme families. While such pooling reduces positional resolution and may obscure localized catalytic effects, it provides a stable and tractable representation for large-scale training. Importantly, RealKcat’s sensitivity to catalytic mutations does not arise from explicit residue-level encoding, but from the curated inclusion of mutant entries and mechanistically guided negatives within KinHub-27k. This supervision compels even pooled embeddings to learn mutation-aware distinctions, enabling generalization across out-of-distribution anchors, synthetic negatives, and dense mutational landscapes.

Our analyses further demonstrate that mutation sensitivity is a defining feature of RealKcat. Across three complementary tiers—out-of-distribution anchors, in-distribution synthetic negatives, and the dense PafA mutational landscape—the model consistently reproduced mutation-induced movements in kinetic bins and maintained bin-level alignment with experimental outcomes. These results highlight the framework’s ability to capture sequence-residue-function relationships beyond trivial memorization, with particular robustness at annotated catalytic residues. At the same time, the PafA evaluation revealed critical limitations which are common to all tools at the moment. RealKcat occasionally underestimates large functional losses, particularly at disordered or densely connected sites such as G56, L297, or H486, where mutational effects propagate through non-local networks that are not fully encoded in sequence embeddings. These observations suggest that while the current model captures catalytic awareness fairly at residue level, it remains limited in resolving mutation effects mediated by structural disorder, allosteric coupling, or long-range interactions. Future incorporation of structure-derived features—such as residue contact density, dynamic flexibility, or cavity context—may address these shortcomings and extend predictive fidelity in challenging regions. We also believe that release of experimental datasets where multiple substrates are tested for activity with the same enzyme (similar to our PafA dataset) will elevate the accuracy and catalytic awareness of RealKcat (and like tools) to the next echelon. One can imagine using updated versions of RealKcat with additional structural priors for identifying mutations to improve activity of target enzymatic reactions.

The implications of RealKcat extend across multiple domains. In enzyme engineering and synthetic biology, the ability to resolve mutation-induced shifts in catalytic parameters provides a rational guide for pathway optimization, reducing reliance on costly experimental screening. In biomedical research, the mutation-sensitive framework offers a predictive platform for prioritizing variants of uncertain significance, enabling early identification of substitutions likely to impair catalytic activity or alter substrate affinity (*33*). Such capabilities are particularly relevant for metabolic disorders and pharmacogenomics, where subtle shifts in kinetic parameters can have outsized physiological consequences (*34*). In systems biology, RealKcat could support enzyme-constrained and proteome-constrained metabolic models by supplying reliable turnover and affinity estimates in magnitude ranges directly compatible with constraint-based formulations (*35–37*). By narrowing parameter uncertainty, it could reduce heavy reliance on stochastic surrogates and improves the interpretability of flux predictions under varying environmental conditions (*38–40*). Across these applications, the emphasis on biologically meaningful classification ranges ensures that RealKcat provides not only accurate predictions but also outputs that can be directly integrated into experimental and computational workflows.

In conclusion, RealKcat represents a step toward biologically interpretable, mutation-aware prediction of enzyme kinetics. Its robustness arises not from algorithmic novelty but from principled dataset curation, mechanistically informed negatives, and carefully chosen evaluation strategies that reflect biochemical practice. While limitations remain in capturing large mutational effects at structurally complex or disordered regions, the framework already establishes a reliable baseline for both predictive and exploratory applications. By embedding enzyme kinetics into an order-of-magnitude classification paradigm, RealKcat aligns computational inference with the practical resolution of experimental biology, offering a versatile tool for enzyme engineering, disease modeling, and systems-level simulations. Future work integrating structural and dynamic descriptors promises to extend these capabilities further, enabling predictive models that not only track catalytic variation but also anticipate the broader consequences of mutations within the complexity of cellular environments.

### Limitations of the Study

Despite its robustness, RealKcat operates under several constraints that define its current scope. The curated KinHub-27k dataset, while extensive, may be biased toward well-characterized classes such as hydrolases and transferases, with translocases and certain ligases underrepresented (**Fig. 1B**), reflecting broader reporting biases in enzymology rather than modeling artifacts. Substrate embeddings derived from isomeric SMILES capture stereochemistry and functional groups but cannot fully represent conformational flexibility or three-dimensional electrostatics, limiting predictions for stereochemistry-rich or structurally complex molecules. Furthermore, framing kinetics as an order-of-magnitude classification task introduces interpretive strengths and weaknesses: e-accuracy acknowledges the physiological variability of enzyme kinetics, which can fluctuate by ±1 log unit across assays due to differences in pH, ionic strength, and temperature (25, 29–31), yet it cannot resolve subtle intra-bin effects, as seen in the PafA analysis where R164A, R164G, and R164V collapsed into the same kinetic cluster despite differing severities of impairment. In addition, RealKcat shows a conservative fallback toward wild-type predictions when challenged with severe mutations, particularly in out-of-distribution or structurally dense contexts. This bias masks the true magnitude of loss for mutations at loop-like or highly interconnected regions such as G56, L297, and H486. The construction of synthetic negatives through alanine substitution at catalytic residues similarly enforces a binary drop/no-drop labeling scheme that strengthens catalytic sensitivity but risks overstating inactivation and neglecting nuanced partial-function variants, an issue compounded by inconsistent metadata on assay conditions within KinHub-27k. Finally, while the dataset includes about 10,000 mutant entries, it is skewed toward loss-of-function variants, offering limited exposure to gain-of-function or context-dependent activity enhancements. Addressing these limitations will require principled integration of structural priors (e.g., residue depth, cavity accessibility, contact density), incorporation of condition-aware modeling to capture assay-dependent variability, and expansion toward balanced mutational landscapes. Notably, naive structural integration has been shown in related work (e.g., CatPred (*10*)) to degrade performance when context is not carefully modeled, underscoring that future refinements must be guided by mechanistic insights and usage of a catalytically relevant sub-structure for each enzyme that stabilizes the active site through direct hydrogen bonding or allostery. We have put forward a recipe to collate such sub-structures for >19,000 experimentally crystallized hydrolases in a reaction core database (RC-Hydrolase) which is now housed within the Protein Data Bank (*41*). These are purported to be stronger regulators of model performance than data volume.

## Materials and Methods

### Database Building

In this study, enzyme kinetic parameters, specifically *k_Cat_* and *K_M_*, were systematically collected and curated from BRENDA [release 2023_1] (*12*) and SABIO-RK [as of May 2024] (*13*), using custom Python scripts [steps in **Fig. 7**]. Initially, 237,487 entries from BRENDA and 77,663 from SABIO-RK were aggregated, parsed for annotations (UniProt ID, organism name, EC number, substrate name, mutant site information and kinetic values), and filtered to retain entries with non-missing fields including those with unique PubMed IDs (PMIDs) to aid verification downstream. This resulted in 96,782 entries from BRENDA and 54,906 from SABIO-RK. Substrate isomeric SMILES keys were obtained from PubChem database (*42*) using IUPAC name of substrates, ensuring consistency across databases to eliminate redundancies. We specifically used the InChI intermediate rather than directly downloading SMILES strings from PubChem, since multiple non-canonical SMILES notations are often stored for the same compound. Conversion through InChI ensured a unique, standardized representation of each substrate before canonicalization with RDKit, thereby minimizing structural inconsistencies and improving reproducibility across downstream embeddings. After extensive data cleaning, entries from both databases were merged, incorporating sequence data from UniProt (*15*) and BRENDA. Sequences with UniProt IDs were directly fetched, while those without UniProtIDs were obtained by querying BRENDA and UniProt using EC number and organism name, followed by applying mutation annotations to adjust sequences accordingly. This process resulted in a database containing 33,442 *k_Cat_* and 48,103 *K_M_* entries, and a merged dataset of 26,244 entries with both *k_Cat_* and *K_M_* values, including 10,244 mutant entries spanning 2,158 scientific articles with unique PubMed IDs (PMIDs).

**Fig. 7.**
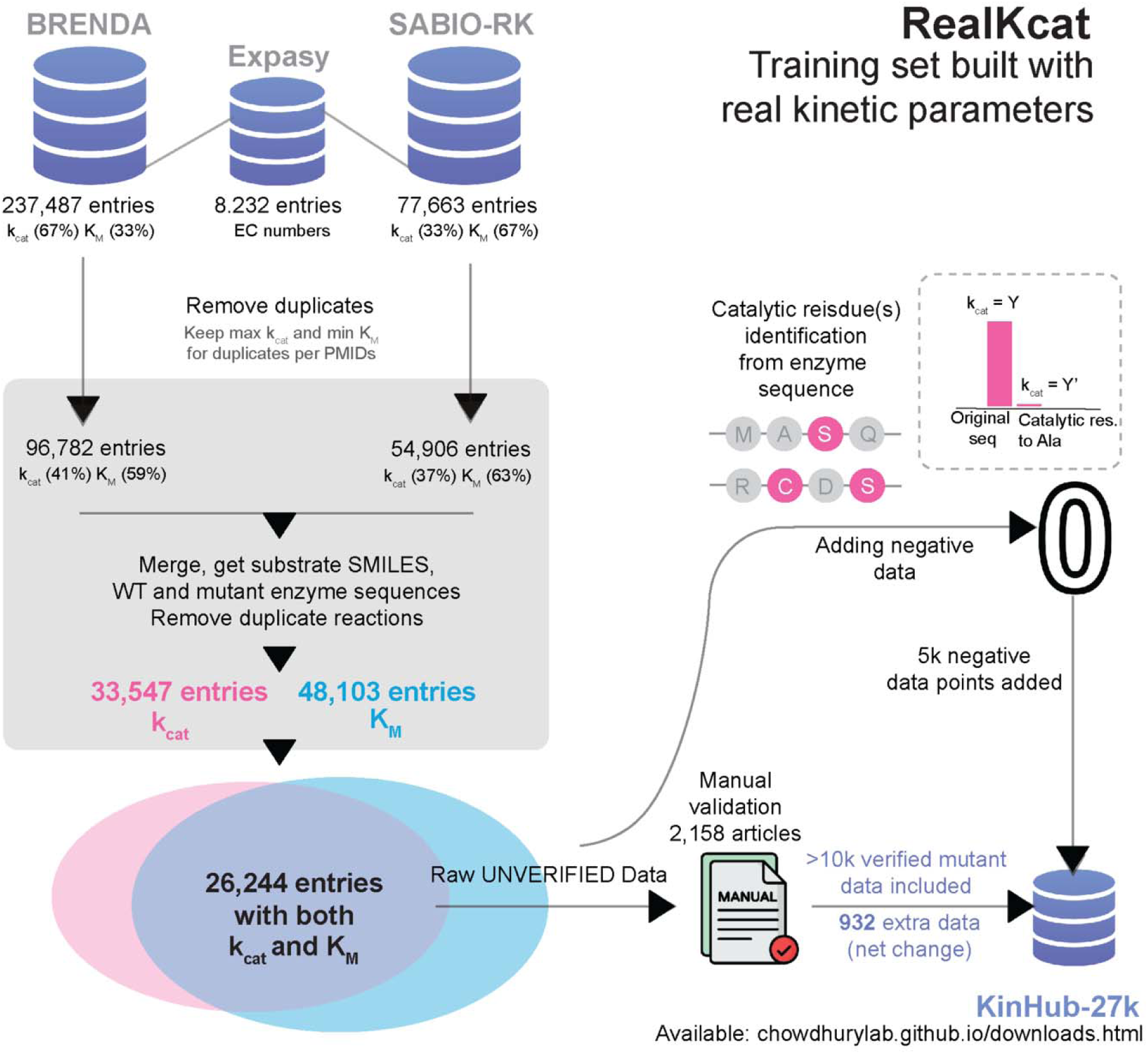
Data collection steps for constructing KinHub-27k from BRENDA and SABIO-RK databases. Raw enzyme kinetic data were sourced from BRENDA (237,487 entries) and SABIO-RK (77,663 entries), covering 8,232 unique EC numbers. Initial processing involved removing duplicate entries while retaining maximum and minimum *K_M_* values, resulting in 96,782 entries from BRENDA and 54,906 from SABIO-RK. After merging the datasets, obtaining SMILES for substrates, and distinguishing between wild-type and mutant sequences, further duplicates were removed, yielding 33,547 unique entries for and 48,103 for *K_M_*. The resulting dataset, KinHub-27k, contains 26,244 entries with paired and *K_M_* values, prepared for manual validation through article review of the original data sources. Note that negative datapoints were added for training but not assigned = 0 (see Materials and Methods for labeling policy).

Following merging, duplicate entries sharing the same isomeric SMILES–sequence pair (i.e., identical enzyme–substrate combinations, regardless of publication source) were collapsed by taking the median value of each kinetic parameter. This refinement replaced the popular maximum *k_Cat_* and minimum *K_M_* filtering approaches, which—while useful for highlighting theoretical performance limits—proved sensitive to atypical or poorly replicated measurements (Fig. S1). Reported values for the same enzyme–substrate pair often varied substantially due to differences in experimental conditions, assay protocols, and the range of organisms or expression systems from which the enzymes were sourced. Median pooling therefore provided a more robust representative value for each unique pair, minimizing bias from outlier measurements while preserving the underlying biological signal across diverse physiological and environmental constraints. The resulting dataset—comprising 14,709 wild-type and 9,808 mutant (MD) entries—captures a broad spectrum of catalytic and affinity ranges while avoiding redundancy.

Prior to median pooling, we do recognize that a significant proportion of these entries might have discrepancies compared to the original references due to data entry errors (*24*). To this end, a manual curation process was conducted. Each article associated with the respective PMIDs for the mutant entries was meticulously reviewed. This rigorous curation process resulted in 11,175 collected mutant entries with *k_Cat_* and *K_M_* values, resolving inconsistencies in 1804 entries and adding 1,072 new entries (Table 2). For instance, the revised *k_Cat_* and *K_M_* values for the human m-NAD-malate dehydrogenase (D102E) mutant in the conversion of fumarate to pyruvate are 60.61 s□¹ and 0.00091 M, respectively—correcting the previously reported erroneous values of 44.14 s□¹ and 0.00255 M. Similarly, for the *Lactobacillus pentosus* D-lactate dehydrogenase (Y52A) mutant catalyzing 2-ketocaproate, the corrected *k_Cat_* and *K_M_* values are 102 s□¹ and 0.0007 M, replacing the inaccurate earlier values of 0.7 s□¹ and 0.102 M. We identified and corrected inaccuracies in reported *K_M_* (788 instances), *k_Cat_* (618 instances), mutants (18 instances), and substrates (240 instances). This curated mutant database, with 16,001 wildtype (WT) and 11,175 mutant (MD) entries with both *k_Cat_* and *K_M_* now forms the curated KinHub-27k dataset that serves as input for preprocessing which then goes into training ML models to predict kinetic parameters.

To capture sequence similarity among enzyme classes, we constructed a network graph. Protein sequences were standardized to a fixed length of 1024 characters through truncation or padding and encoded by their character ordinal values. A fixed length of 1024 characters was used, as ESM-2 or ESM-C does not accept sequences longer than this length. Pairwise sequence identity was computed across all unique enzyme sequences using all-vs-all global alignments (Biopython pairwise2.align.globalxx, implementing the Needleman–Wunsch algorithm with unit match scores and zero mismatch/gap penalties). The resulting percent identity scores were used to construct a graph representation, with distances defined as 1−identity. A minimum spanning tree (MST) was extracted to provide a parsimonious backbone of the sequence space, and each node was annotated by its primary EC class. The network was visualized using a distance-aware layout (**Fig. 1C**), thereby highlighting intra-class homology while also capturing the global diversity of sequence relationships represented in KinHub-27k.

### Binning Strategy for Enzyme Kinetic Parameters

In RealKcat, continuous enzyme kinetic parameters *k_Cat_* and *K_M_* values are reformulated as multi-class classification tasks to address the substantial order-of-magnitude variability inherent in enzyme kinetics and the practical limitations of direct regression. Kinetic parameters were then discretized into biologically meaningful, log_10_-spaced bins, enabling robust range prediction while reducing the impact of experimental noise. Specifically, this binning process categorizes *k_Cat_* values into ranges, each representing an order of magnitude difference, while reserving specific clusters for extreme values (refer to Figure 1F, 1G, and 1H). This binning approach involved defining specific cluster edges: *k_Cat_* values were categorized with boundaries set at [0,10^−8^,10^−2^,10^−1^,10^0^,10^1^,10^2^,10^3^,10^8^] s^−1^ and *K_M_* values at [10^−14^,10^−5^,10^−4^,10^−3^,10^−2^,10^−1^,10^4^] M. The classification approach provides a structured prediction strategy, allowing the model to interpret parameter ranges with high fidelity and minimizing the impact of noise from imprecise continuous predictions.

### Mechanistically Guided Synthetic Negatives and Hybrid Relabeling

To model catalytic loss events and expand training coverage, we constructed a mechanistically guided synthetic negative dataset from UniProt-annotated catalytic, substrate-binding, and cofactor-binding residues. Each wild-type sequence underwent alanine-scanning mutagenesis, a strategy that preserves backbone conformation while removing side-chain chemistry to selectively attenuate catalytic function with minimal structural disruption. For multi-residue active sites, all annotated active site residues were substituted simultaneously to emulate severe functional impairment. This process yielded 5,278 high-confidence synthetic negatives, including 22 experimentally curated wild-type–mutant (WT–MD_Ala_) pairs spanning diverse enzyme–substrate contexts (see Supplemental Table 1). A ≤60% global sequence identity threshold was enforced only between all data subsets and the 22-sample synthetic negative anchor set to maintain OOD separation. The similarity distributions shown in Fig. 3D–E were computed between the test and training partitions for performance stratification and may therefore include identities exceeding 60%. Kinetic effects in these curated pairs were quantified as the bin difference *Δb=b_WT_−b_MD_* on the clustered kinetic scale (i.e. the same log_10_-spaced bins employed in the data preprocessing pipeline). For *k_Cat_*, a significant drop was defined as *Δk_Cat_ ≥*2 bins (i.e. ≥ two orders-of-magnitude reduction), ensuring that only major catalytic losses were labeled positive. For *K_M_*, a drop was defined as Δ*K_M_* ≤−1 bin (≥ one order-of-magnitude increase), reflecting weaker substrate affinity. This asymmetric thresholding is consistent with biochemical expectations that active-site disruptions predominantly affect turnover rather than binding, unless direct contact residues are altered.

Because most synthetic negatives lacked experimental kinetic measurements, we implemented a two-stage relabeling framework. The curated WT–MD_Ala_ pairs served as the training set for a meta-learned binary classifier, which was then used to assign **drop/no-drop** labels to the broader unlabeled synthetic set. For each enzyme–substrate pair, we computed a suite of mechanistically relevant descriptors spanning mutational severity, substrate chemistry, and interaction scale. Mutational severity was quantified as the sum of BLOSUM62 substitution scores for alanine replacement of annotated residues, providing an evolutionary substitution cost metric. Substrate chemistry was characterized by molecular weight, Crippen’s logP, total ring count, hydrogen bond donor count, and hydrogen bond acceptor count from RDKit-parsed isomeric SMILES. Interaction-scale metrics included active-site density (annotated residues per sequence length) and substrate-to-enzyme molecular weight ratio, with the wild-type kinetic bin for the relevant parameter included as a baseline activity indicator.

Binary classification for both *k_Cat_* and *K_M_* drop prediction tasks was performed using the *AutoTabPFNClassifier*, a Transformer-based meta-learned prior optimized for tabular data (*43*). Given the small curated set, Leave-One-Out Cross-Validation (LOOCV) was employed to maximize data usage while maintaining strict independence between training and test samples. In each fold, features were standardized using a *StandardScaler* fit exclusively on the training set to prevent information leakage. *AutoTabPFN* models were trained with a per-fold time limit of 1000 seconds on GPU and evaluated via classification accuracy and aggregated confusion matrices. Final models for *k_Cat_* and *K_M_* were retrained on their complete datasets and serialized alongside their scalers using *joblib*, enabling reproducibility and deployment within the downstream RealKcat training pipeline. Hence, the resulting binary classifiers were used to annotate unlabeled synthetic negatives as drop or no-drop. These labels informed *a hybrid relabeling policy*: in *k_Cat_* tasks, drops were randomly reassigned to bins well below (≥ two orders-of-magnitude reduction) the WT value and non-drops constrained within ±1 bin; in *K_M_* tasks, drops shifted to higher bins (≥ one order-of-magnitude increase) while non-drops were restricted to WT or lower bins. Thus, ensuring kinetic plausibility, generating hard negatives that are sequence-close to actives but mechanistically divergent in catalytic space.

### Partitioning Strategy for Leakage-Minimized Data Splits

To prevent over-representation of densely sampled regions and maintain structural diversity, the negative dataset was partitioned using a manifold-guided strategy. The enzyme subspace (the first portion of the concatenated features corresponds to enzyme embeddings) was projected into two dimensions using PaCMAP, a dimensionality reduction method that preserves both global and local relationships while mitigating distortions common in high-dimensional Euclidean space. Affinity Propagation (AP) clustering was then applied to the PaCMAP projection to identify exemplar-based sequence clusters. Clusters containing fewer than five members were reassigned to a “miscellaneous” bin (label −1) to prevent small groups from disproportionately influencing stratification. Partitioning proceeded in two stages. First, a stratified hold-out split with respect to AP cluster labels and their associated drop/no-drop classifications for both *k_Cat_* and *K_M_* set aside 5% of the negative set as a temporary pool, preserving the empirical distribution of these labels (including the −1 “miscellaneous” bin). Second, the temporary pool was bisected evenly into validation and test subsets, yielding approximate final proportions of ∼95%/2.5%/2.5% for train/val/test. All random seeds were fixed (random_state=42) to allow exact regeneration of splits across runs.

To preserve biological representativeness and statistical rigor, we implemented a two-tier, source-aware positive set partitioning framework. The aim was to maintain the endogenous balance between wild-type (WT) and mutant-derived (MD) entries while controlling for potential covariates between biological source and kinetic class. First, a catalytic inference hold-out of 100 positive samples was constructed by proportionally sampling from WT and MD pools without replacement, preserving the natural WT:MD (60:40) ratio and ensuring statistical independence from the training data. This was performed under a fixed random seed of 42. In parallel, a negative hold-out of 10 synthetic variants was selected at random from the partitioned negative test pool. These negatives were generated by alanine substitution at annotated catalytic residues. Although their corresponding wild-type sequences are present in the positive training partitions, the exact negative sequences were excluded from training. Consequently, this configuration ensures that negatives are evaluated in-distribution with respect to enzyme identity but remain unseen as explicit negative examples. This two-tier scheme enables evaluation across both positive (WT and MD) and synthetic negative chemotypes, providing a controlled framework for assessing generalization and mutation sensitivity.

The remaining positives underwent five-fold cross-validation stratified on a composite label that jointly encoded kinetic bin and biological source (WT/MD). This ensured balanced representation of both factors in every fold. Negative sets were handled in a paired, contrastive manner: synthetic negatives corresponding to a given WT enzyme were co-assigned to the same partition as their cognate positives. This pairing preserved the biological relevance of evaluation and ensured that synthetic negatives were consistently matched with their related WT samples across train/validation/test splits. Within each fold, the outer 20% of the data was evenly divided into validation and test subsets under the same composite-label stratification. This procedure yielded approximate 80/10/10 proportions for train/validation/test in each fold, while preserving both kinetic class distribution and WT:MD balance. In addition, all partitioning steps—including manifold-guided clustering for the negative set, proportional WT:MD sampling for the positive set, and positive–negative pairing—were performed with a fixed random seed (42). This ensured that every split could be exactly reproduced, maintaining consistency across experiments and enabling rigorous model comparison.

### Feature Representation and Embeddings

In RealKcat’s feature extraction pipeline, each sample from the curated KinHub-27k dataset comprises an enzyme sequence, an isomeric SMILES representation of the substrate, and assigned log-binned clusters for *k_Cat_* and *K_M_*. Enzyme sequences were first evaluated using embeddings from both ESM-2 (esm2_t33_650M_UR50D) and ESM-C (600M; EvolutionaryScale/esmc-600m-2024-12), protein language models trained on large-scale sequence corpora to produce context-aware embeddings (*11*, *25*). For both models, the CLS ([0]) and terminal/padding tokens were excluded, and residue-level embeddings across positions 1…L (where L is sequence length) were mean-pooled to produce sequence descriptors, ensuring that representations reflected residue content rather than global classification tokens. Embeddings were drawn from the final layers (layer 33 for ESM-2 and layer 36 for ESM-C) to capture conserved evolutionary signals, sequence-context dependencies, and biochemical constraints across enzyme families.

Although mean pooling inevitably reduces positional resolution and may obscure localized effects at catalytic residues, it provides a stable and computationally tractable representation for large-scale training. As our evaluations showed, this approach retains sufficient sensitivity to mutation-induced perturbations when paired with rigorous curation and negative sampling. Between the two models, both ESM-2 (1,280-dimensional) and ESM-C (1,152-dimensional) supported robust performance, but ESM-C consistently offered modest advantages in mutation-sensitive and out-of-distribution tests (see Results, **Fig. 3**). Based on these comparative results, ESM-C was adopted as the default enzyme representation in RealKcat.

On the other hand, substrate features were encoded using **ChemBERTa** (seyonec/PubChem10M_SMILES_BPE_450k) embeddings, derived from isomeric SMILES strings and averaged across all tokens (including the CLS token) including the CLS token of final hidden states to yield 768-dimensional molecular descriptors of all subtrates/co-factor/ions. This whole-sequence pooling preserves information from both token-level chemical substructures and the global sequence summary embedded in the CLS representation, capturing stereochemistry, functional groups, and physicochemical properties directly relevant to substrate recognition and catalysis (*18*).

We note that CLS token handling differed between enzyme and substrate embeddings. For ChemBERTa, we retained the CLS token in the mean-pooling operation, since this embedding has been pretrained to act as a global sequence summary that captures molecular-level context beyond token-local features. In contrast, for ESM protein embeddings, the CLS token was excluded, as prior benchmarks indicate that mean pooling across residue tokens (excluding CLS and padding) yields more stable and biochemically relevant sequence descriptors. This design choice reflects the distinct pretraining objectives of the two models and was confirmed in our diagnostics to provide more consistent performance across tasks.

For each enzyme–substrate pair, the ESM-C (1,152-D) and ChemBERTa (768-D) embeddings were concatenated to form a unified 1,920-dimensional joint feature vector, integrating both evolutionary context and substrate chemistry in a balanced representation. This formulation does not enforce explicit interaction modeling, but rather provides a feature space from which the classifier can learn associations between catalytic variation and molecular context. Prior to model training, all features were globally standardized, and partition-specific normalization parameters were estimated strictly on training folds to avoid leakage.

### Model Architecture and Training

The architecture of the RealKcat model centers on a highly optimized gradient-boosting framework, specifically utilizing the Extreme Gradient Boosting (XGBoost) algorithm. XGBoost is a robust machine learning model, particularly well-suited for handling high-dimensional, non-linear data and yielding high accuracy in multi-class classification contexts (*26*), which aligns with the complexities of enzyme kinetics prediction. In RealKcat, XGBoost constructs an ensemble of decision trees, where each tree 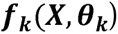 is sequentially added to the model, aiming to minimize the residual errors left by the previous trees. This process optimizes a compound objective function: 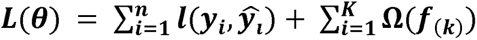 where 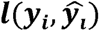 denotes the log-loss for multi-class classification, specifically capturing the errors between true labels ***y_i_*** and predictions ***ŷ_i_***. The second term, 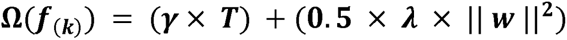, is a regularization penalty that controls model complexity by penalizing each tree’s structural complexity, with ***T*** representing the number of leaves in a tree and ***w*** the leaf weights. This regularization ensures that the model balances the trade-offs between bias and variance, which is crucial in kinetic prediction tasks where enzyme-substrate interactions are diverse. ***γ*** (**gamma**) parameter controls the structural complexity of each tree by penalizing the addition of new leaves. Specifically, **γ** determines the minimum gain a node must produce to be further split. Higher values of ***γ*** make the model more conservative, favoring trees with fewer leaves by only allowing splits that significantly improve the model. This prevents the formation of overly detailed trees, which could lead to overfitting. However, increasing **T** can lead to overfitting, as it allows the tree to become more tailored to the training set. The term **γ** × **T** thus penalizes trees with large numbers of leaves, encouraging simpler structures that are less likely to overfit.

Defining other components: **λ (lambda)**: This is the **L2 regularization parameter** on leaf weights, and it helps to control the size of the weights associated with each leaf. A higher value of **λ** increases the penalty on large weights, which discourages the model from relying too heavily on any single leaf’s prediction. By controlling the magnitude of leaf weights, **λ** helps improve the model’s robustness, making it less sensitive to variations in the data. || ***w*** ||**^2^**: This term represents the **squared L2 norm of the leaf weights**, where ***w*** is a vector containing the prediction values at each leaf node in the tree. The squared norm || ***w*** ||**^2^** calculates the sum of the squares of these weights, which quantifies the overall “strength” of the predictions at the leaves. By penalizing larger norms, the regularization term helps to distribute the prediction values more evenly across leaves, thus reducing the likelihood of overfitting. **0.5** × **λ** × || ***w*** ||**^2^**: This part of the regularization term penalizes large leaf weights while controlling the model’s complexity by discouraging the creation of overly powerful leaves. The factor of 0.5 is used to scale this penalty, allowing XGBoost to balance the model’s accuracy with its simplicity more effectively. Together, these components of 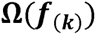 provide nuanced control over the model’s complexity. By adjusting **γ** and **λ**, the user can fine-tune how aggressively the model penalizes complexity in terms of both the number of leaves and the magnitude of leaf predictions, ultimately promoting a balance between model accuracy and generalizability.

XGBoost further improves predictive accuracy by utilizing gradient-based ***pseudo -residuals***, computed as the gradient of the loss function with respect to current predictions, thereby emphasizing instances with high residual error from previous rounds. This refinement process, regulated by a learning rate ***η***, enables RealKcat to iteratively correct errors while stabilizing updates, effectively capturing subtle variations in kinetic parameters across different enzyme classes. Mathematically, the **additive** update for each tree in the boosting process can be described by: 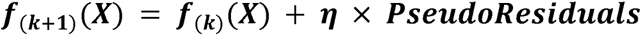, where ***η*** is tuned to gradually refine predictions without over-adjusting for noise in the data.

#### Training and Hyperparameter Optimization

Model hyperparameters were optimized through a random-search strategy implemented in Optuna under the earlier described non-conventional stratified 5-fold cross-validation scheme. Stratification preserved the distribution of kinetic parameter bins across folds, ensuring that both abundant and minority classes were adequately represented in each split. The Optuna search optimized over validation accuracy as the objective function, with candidate models evaluated on held-out validation partitions to prevent information leakage. The final hyperparameters selected for downstream training were: number of estimators = 1920, maximum depth = 11, learning rate (η) = 0.0021, maximum delta step (δ) = 0, and L1 regularization (α) = 1.13, L2 regularization (λ) = 14.803. Early stopping was employed with a patience of 10 rounds on the validation negative log-loss (cross-entropy loss) as the monitoring metric, providing an effective safeguard against overfitting while preserving generalization.

To address class imbalance, particularly in underrepresented kinetic bins, Synthetic Minority Over-sampling Technique (SMOTE) oversampling was applied exclusively to the training partition of each fold following the method of Chawla *et al.* (*28*), thereby enhancing representation without leakage into validation or test sets. SMOTE generates synthetic minority-class instances by interpolating between a given sample and its k-nearest neighbors, thereby enhancing representation without altering class labels. We used the default parameter *k=5*, which yielded an approximately even distribution across kinetic bins while preserving intra-class variability. For reproducibility, a fixed random seed (42) was applied to ensure identical synthetic instance generation across runs. Importantly, oversampling was performed **after binning**, allowing targeted correction within each order-of-magnitude kinetic bin—an essential step given the naturally skewed distribution of enzyme kinetics data.

Synthetic negatives, introduced via the hybrid relabeling policy, were further down-weighted by a constant factor (0.001) to reduce their disproportionate influence while retaining their calibration role; weights for positive entries were left unchanged. All weights were assigned in full-batch form, and datasets were shuffled with a fixed random seed (42) for reproducibility. Following the initial training pass, an optional hard negative mining (HNM) procedure was applied: misclassified training examples—especially false negatives not within ±1 order of magnitude —were identified, their sample weights boosted by a factor of 10^2^, and the reweighted examples reintroduced into the training pool. The model was then fine-tuned for an additional ≥10 boosting rounds (approximately 20% of the original boosting iterations), starting from the previous booster state. This two-stage scheme improved sensitivity to difficult decision boundaries, particularly for mutation-sensitive bins and structurally challenging negatives, while maintaining stability and interpretability across cross-validation folds.

Moreover, in our 5-fold stratified cross-validation scheme, each fold produced an independently trained RealKcat model with its own training/validation/test split. For benchmarking, we reported mean ± SD performance across all five folds (to reflect population-level reproducibility), but for downstream inference we needed a single deployed model. To this end, we selected **fold-5** as the representative model for inference. Fold-5 was chosen a priori to be held aside as the final inference model in our experimental design, avoiding any post-hoc cherry-picking.

#### Feature Standardization

Given the heterogeneous nature of the input features, each embedding group—ESM and ChemBERTa—was independently standardized using the mean and standard deviation from the training set. Specifically, each feature ***x*** in the embedding was transformed as follows: 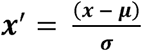, where ***µ*** and ***σ*** are the mean and standard deviation of the feature in the training set. This standardization was applied consistently across validation and test sets, ensuring a stable feature distribution across datasets, aiding in model convergence, and enhancing generalization across diverse kinetic parameter classes

### Performance and Evaluation Metrics

RealKcat’s predictive capacity was evaluated using a suite of metrics specifically chosen for multi-class classification under class imbalance. Standard accuracy, 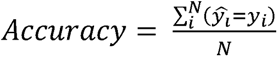, defined as the proportion of exact bin matches between predicted (*ŷ_i_*) and true (*y_i_*) labels, provides a baseline measure of overall correctness but is limited in this context due to the skewed distribution of kinetic parameters across bins. To complement this, we reported class-specific precision, recall, and F1-scores, macro-averaged across classes to prevent dominance by majority bins.

A central metric in this study is the **error-tolerant accuracy (e-accuracy)**, which defines correctness within ±1 log_10_-spaced bin of the experimental label: 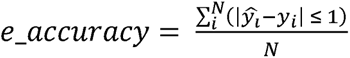, reflecting the biological reality that order-of-magnitude deviations rarely alter functional interpretation. To further assess balanced classification, we used the **Matthews Correlation Coefficient (MCC),** macro-averaged across bins, which incorporates all entries of the confusion matrix and is less sensitive to class imbalance than raw accuracy. Finally, the **area under the precision–recall curve (AUC-PR)** was included as it provides a more informative measure than ROC curves under imbalanced conditions.

Visual diagnostics complemented these quantitative measures. **Confusion matrices** captured per-bin prediction fidelity, highlighting both central and extreme kinetic ranges, while **t-SNE projections** of concatenated embeddings overlaid with predicted versus true classes illustrated preservation of the global kinetic parameter space. All metrics, including accuracy, precision, recall, F1-score, e-accuracy, MCC, and AUC-PR, were implemented in Python using the *scikit-learn* (sklearn) library.

### Mutation-Aware Validation Datasets

To assess mutation sensitivity, we curated three complementary datasets. The first comprised 22 experimentally characterized wild-type and alanine-substituted pairs at UniProt-annotated catalytic residues, withheld entirely from training and used as an out-of-distribution (OOD) anchor set. The second comprised a synthetic hold-out of 10 alanine-substituted variants randomly drawn from the partitioned negative pool; their corresponding wild-type sequences were included in the positive training set, but the negative variants themselves were withheld, yielding an in-distribution-by-identity benchmark. For both sets, mutation effects were additionally summarized using binary meta-labels (*k_Cat_* drop vs no-drop, *K_M_* increase vs no-increase) derived from mechanistic descriptors. An additional set of 10 non-alanine OOD mutants (≤ 80 % sequence identity relative to the training partition) was further used to confirm that performance generalizes beyond alanine substitutions (see Fig. S4). Finally, we employed the high-throughput alkaline phosphatase PafA from *Elizabethkingia meningoseptica* mutational dataset, consisting of 1,016 single-point variants assayed with carboxy 4-methylumbelliferyl phosphate ester (cMUP) substrate (*29*). This dataset was partitioned into 398 training, 310 validation, and 310 test variants, with WT and one catalytic variant (T79S) included in training, while catalytic-site substitutions at residue R164 (R164A, R164G, R164V) were withheld in the test/validation sets. Performance was quantified using e-accuracy, MCC, and meta-label classification metrics as detailed under *Performance and Evaluation Metrics*.

### Computational Resources

This study leveraged the high-performance computational resources of the University of Nebraska-Lincoln’s SWAN HCC cluster, operating on Ubuntu Linux. Our primary system setup featured dual Intel(R) Xeon(R) Gold 6248R CPUs, totaling 48 cores (used only 16 cores) and 187 GiB of RAM, of which approximately 143 GiB was available for handling large datasets. For GPU-accelerated tasks, we employed an NVIDIA Tesla V100S-PCIE-32GB, running CUDA 12.4 with NVIDIA driver version 550.127.05, which provided 32 GiB of VRAM—around 17.9 GiB of which was dedicated to embeddings and deep learning computations. Additionally, the Iowa State University high-performance computer NOVA contributed to data curation, embeddings, and model training, utilizing dual 18-core Intel Xeon Gold 6140 CPUs (36 cores total), 192 GiB RAM, and dual NVIDIA Tesla V100-32GB GPUs with CUDA 11.8 for GPU-accelerated processing.

Several components of this implementation were managed using Python 3.12.5, integrating a robust suite of libraries for machine learning and biochemical data processing. The primary libraries included XGBoost (version 2.1.1) for model training and prediction, and PyTorch (version 2.4.0) for handling tensor operations and neural network computations. To balance class distributions, we used the imbalanced-learn library (version 0.12.3) for SMOTE applications, while Scikit-learn (version 1.5.1) provided tools for calculating evaluation metrics and model performance. For reproducibility, we set a random seed of 42 across all stochastic processes, ensuring consistency in SMOTE applications, train-validation-test splits, and model initialization.

#### Code availability

An open-source notebook with code is made available in a public repository to foster use easy, through the link: https://github.com/TKAI-LAB-Mali/RealKcat. Additionally, we are committed to making all scripts, notebooks, and relevant code available through a public repository ***upon publication***, supporting reproducibility and enabling further exploration by the research community.

## Acknowledgments

A.O. wishes to express gratitude for the financial support provided for this research. This includes the NIH MIRA Award (5R35GM143009) awarded to R.S. This work is also partially supported by the Iowa State University Startup Grant, and NSF EPSCoR RII Track-1, Award Number DQDBM7FGJPC5 to R.C., and Iowa Economic Development Authority Award Number: 24IEC006 to R.C. and R.S.

## Author contributions

Conceptualization: R.C., R.S., A.M., A.O., K.A.S.

Methodology: R.C., A.O., K.A.S., A.M., A.B., S.F.

Investigation: K.A.S., A.O., A.M., A.B., S.F., N.S., M.S.N., S.K., L.M.S., R.S., N.B.C., R.A., B.C.

Visualization: A.O., A.M., M.S.N., S.F., R.C.

Funding acquisition: R.C., R.S.

Project administration: R.C.

Supervision: R.C., R.S.

Writing – original draft: A.O., K.A.S, A.M., A.B.

Writing – review & editing: K.A.S., A.O., A.M., A.B., R.S., R.C.

## Data and materials availability

The entire KinHub-27k dataset and accompanying code is available for download and inference at the link: https://chowdhurylab.github.io/downloads.html. Moreover, an open-source notebook with code is made available in a public repository to foster use easy, through the link: https://github.com/TKAI-LAB-Mali/RealKcat. Additionally, we are committed to making all scripts, notebooks, and relevant code available through a public repository *upon publication*, supporting reproducibility and enabling further exploration by the research community.

