## Supplemental Information for "Robust Prediction of Enzyme Variant Kinetics with RealKcat"

**The PDF file includes:**

Supplementary Text

Figs. S1 to S4

Tables S1

### Supplementary Text

#### ***Justification for e-Accuracy in RealKcat***

While **accuracy** in RealKcat is defined as the percentage of predictions that fall within the exact cluster as the experimental value, **e-accuracy** is defined as the model’s ability to predict enzyme kinetics within one cluster above or below the true experimental cluster. For instance, if the actual 𝑘_𝑐𝑎𝑡_ value falls within cluster **b**, which ranges from 10^1^ to 10^2^, an e-accurate prediction would place the predicted value within either cluster **a** (10^0^ to 10^1^) or cluster **c** (10^2^ to 10^3^). This measure of accuracy provides a practical assessment of the model’s capability to capture functionally relevant ranges, as it acknowledges that slight deviations in the predicted cluster are still valuable for applications where an approximate range suffices.

To illustrate why a robust metric like e-accuracy is important, **Supplementary Fig. S1** shows the observed spread of experimental 𝑘_𝑐𝑎𝑡_ and *K_M_* values for identical sequence–substrate pairs before median pooling. In many cases, values span **6–8 orders of magnitude**, reflecting variation due to differences in experimental conditions, assay protocols, or organism sources. Such large spreads mean that extreme-value (max–min) filtering—used in some prior works—can disproportionately emphasize atypical or poorly replicated measurements, potentially distorting downstream model training and evaluation. By contrast, median pooling yields a stable central estimate for each unique pair, and e-accuracy naturally accommodates the residual spread that still exists after curation by rewarding predictions that land within an adjacent, functionally similar kinetic range. This combination improves robustness in enzyme design and metabolic modeling applications, where capturing the correct order of magnitude is often more critical than pinpointing the exact experimental value.

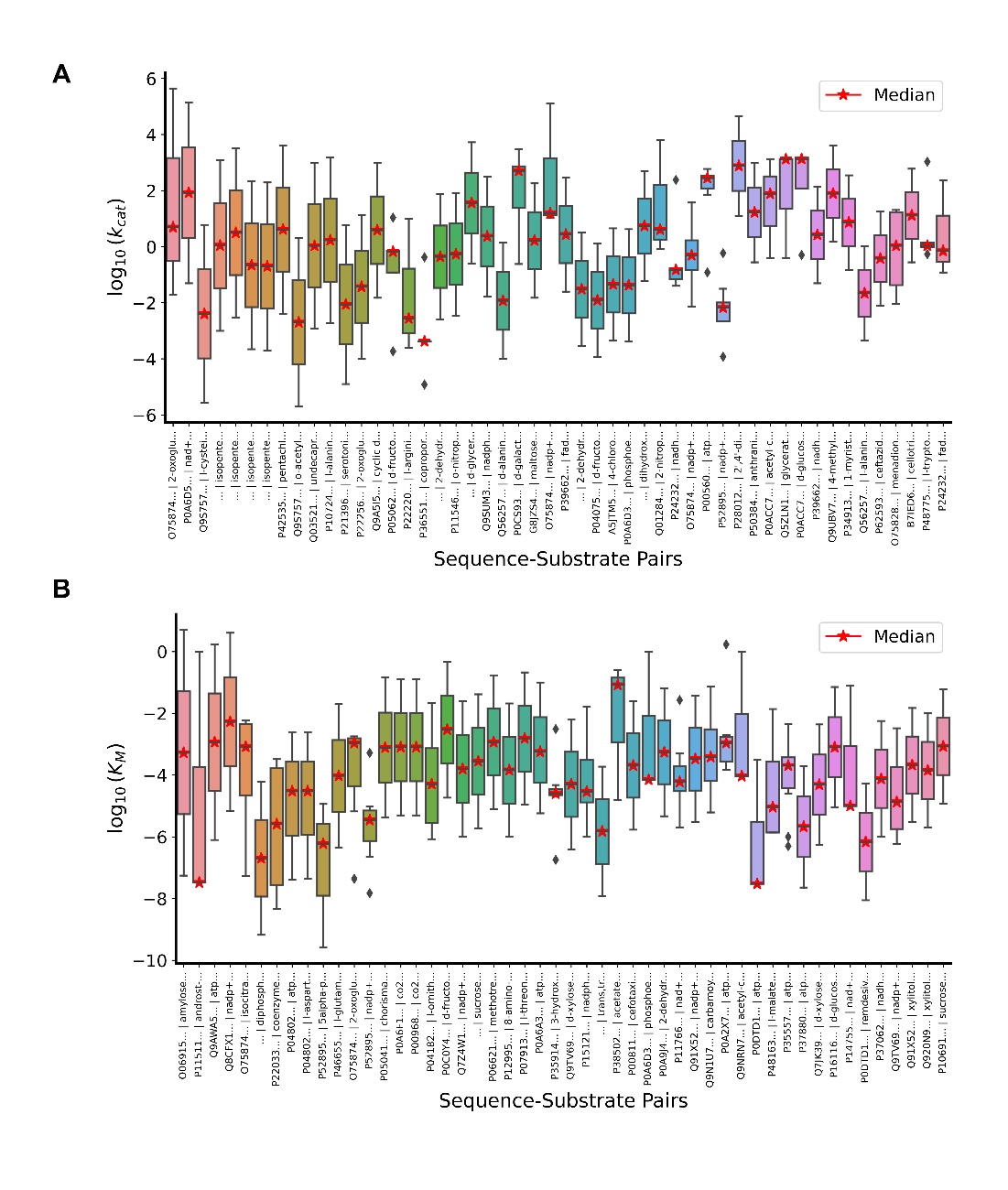

**Fig. S1. Large experimental variation in kinetic parameters for the top 50 enzyme–substrate pairs.**
Boxplots show the distribution of experimental **(A)** 𝑘_𝑐𝑎𝑡_ and **(B)** *K_M_* values for the **50 sequence–isomeric SMILES pairs** with the largest order-of-magnitude spread in reported measurements, aggregated from BRENDA and SABIO-RK. Each box represents all reported values for a given pair, with the red star indicating the median. In many cases, values span **6–8 orders of magnitude**, reflecting variation introduced by differences in assay conditions, measurement protocols, and organism sources. Such wide spreads highlight the limitations of extreme-value (max–min) filtering, which can overemphasize atypical measurements, and motivate the use of median pooling to obtain robust central estimates for downstream modeling.

#### Model Evaluation

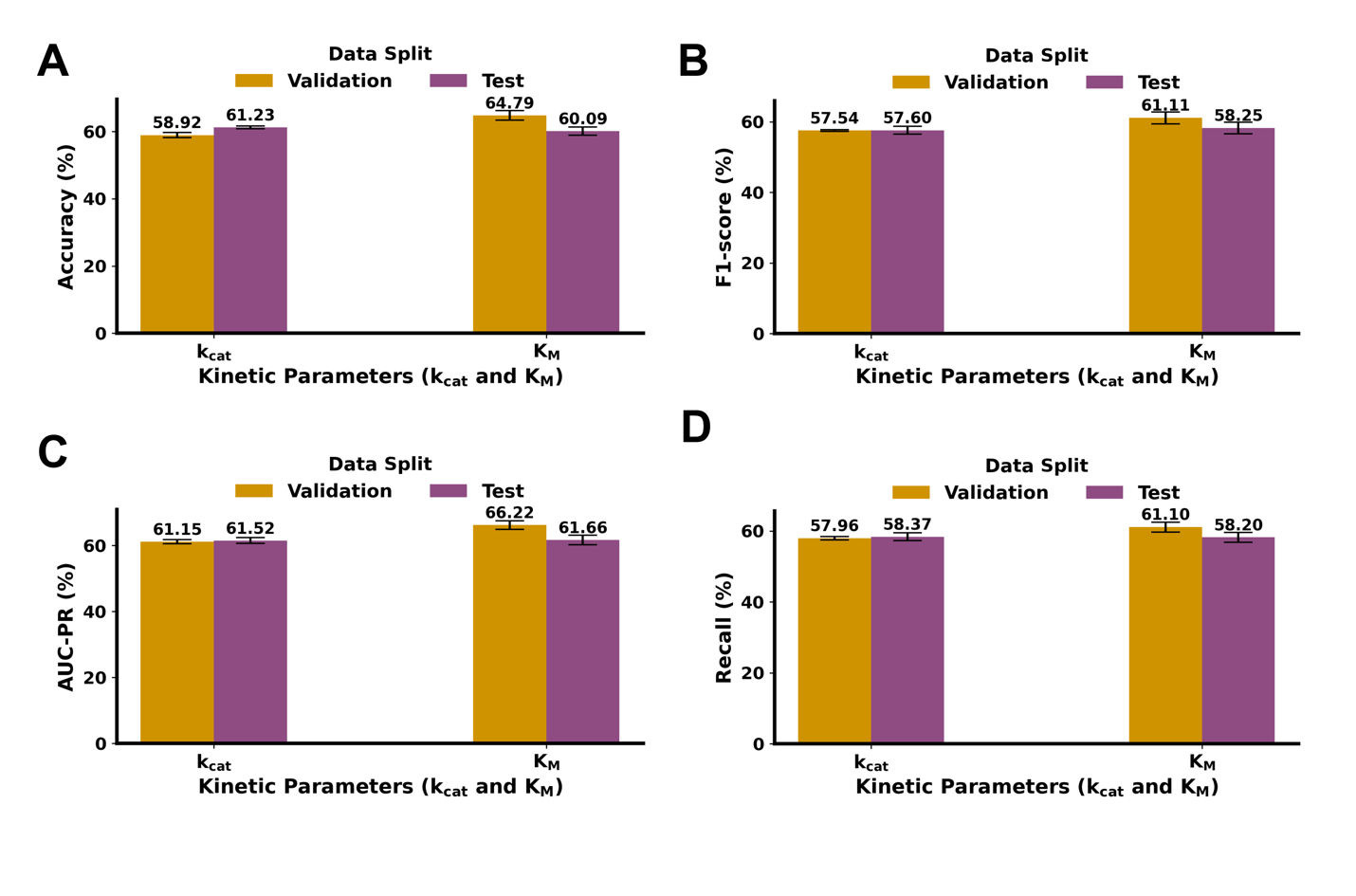

**Fig. S2. Standard performance metrics of RealKcat across data splits for 𝑘_𝑐𝑎𝑡_ and *K_M_*. (A)** Exact-bin accuracy across validation and test sets. **(B)** F1-scores show similar trends, with reduced stability in under-represented classes. **(C)** Area under the precision–recall curve (AUC-PR) and **(D)** recall further illustrate the imbalance-driven variation, with metrics skewed toward majority classes. Panels A–D report the mean across all five folds ± standard deviation. These results underscore why exact-bin accuracy alone may overstate predictive quality in multi-class enzyme kinetics classification. By contrast, the use of **macro-averaged MCC** (reported in Fig. 3) provides a more balanced and conservative measure of performance, as it accounts for all entries in the confusion matrix and reduces the dominance of central clusters. Together with e-accuracy, MCC offers a biologically meaningful and robust evaluation of RealKcat’s ability to generalize across both frequent and rare kinetic regimes.

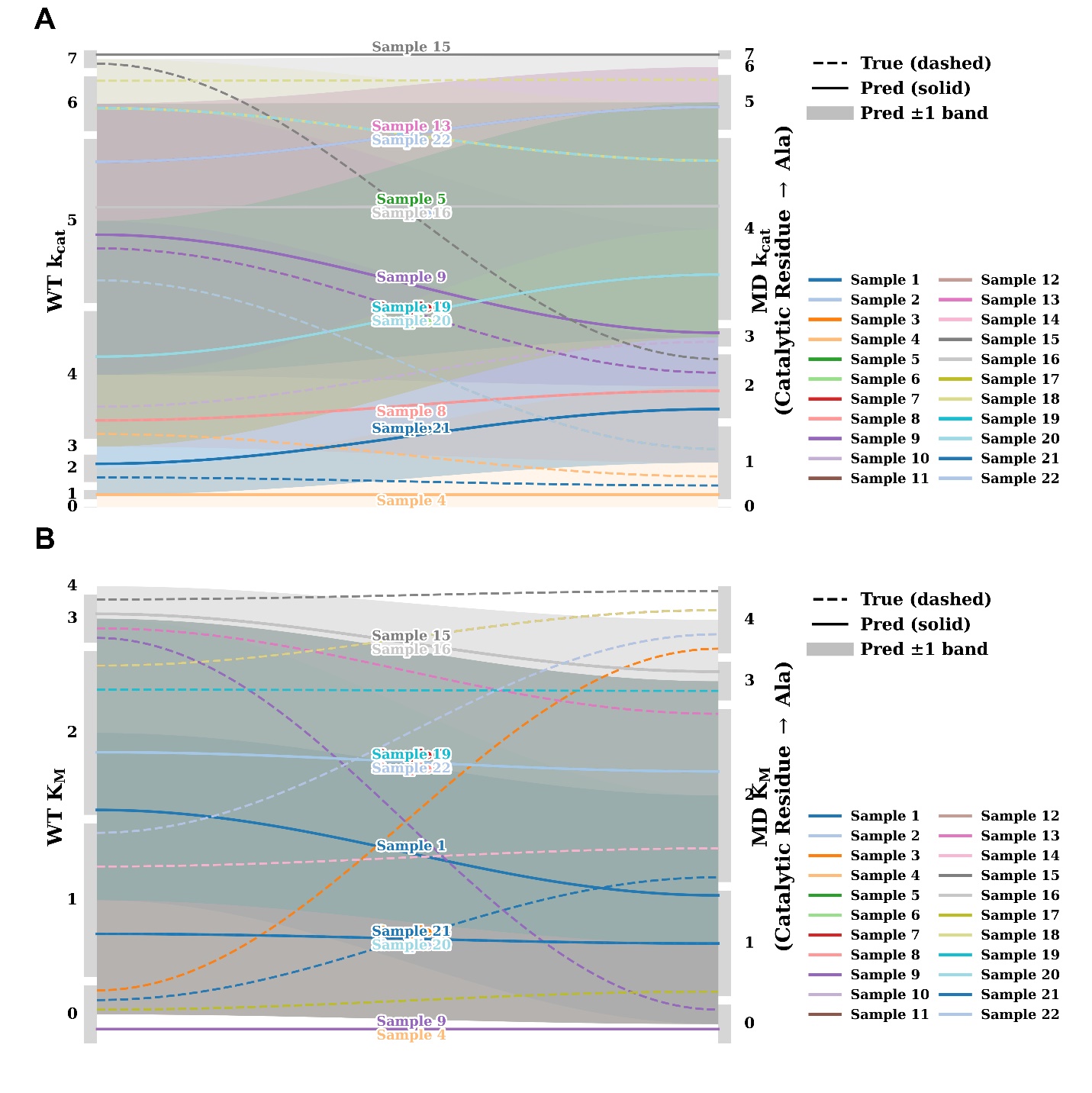

**Fig. S3. Longitudinal visualization of WT–MD*_Ala_* anchor pairs withheld from training.** **(A)** Predicted versus experimental trajectories for 𝑘_𝑐𝑎𝑡_ across 22 WT–MD*_Ala_* pairs. Each colored curve represents a unique enzyme–substrate pair, with dashed lines denoting experimental values and solid lines denoting RealKcat predictions. Grey shading indicates the ±1 bin tolerance window used in e-accuracy assessment. Predicted shifts generally track the experimental trajectories, with most mutant values remaining within the ±1 tolerance band. **(B)** Equivalent analysis for *K_M_* across the same 22 WT–MD*_Ala_* pairs. Again, RealKcat predictions (solid lines) reproduce the directionality of experimental changes (dashed lines), with bin-level agreement preserved in the majority of cases. This complementary visualization highlights that RealKcat maintains the relative ordering of catalytic parameters across WT–mutant pairs, further supporting the results in **Fig. 4**.

**Table S1. Curated set of 22 experimentally validated wild-type (WT)–mutant alanine (MD*_Ala_*) pairs at UniProt-annotated catalytic residues, used to anchor the mechanistically guided negative dataset.** These entries consistently exhibit substantial catalytic impairment—typically ≥2 orders-of-magnitude reductions in 𝑘_𝑐𝑎𝑡_ and ≥1 order-of-magnitude increases in *K_M_*—providing benchmarks for calibrating synthetic negatives and for training the binary meta-label classifier.

Table_S1.xlsx attached

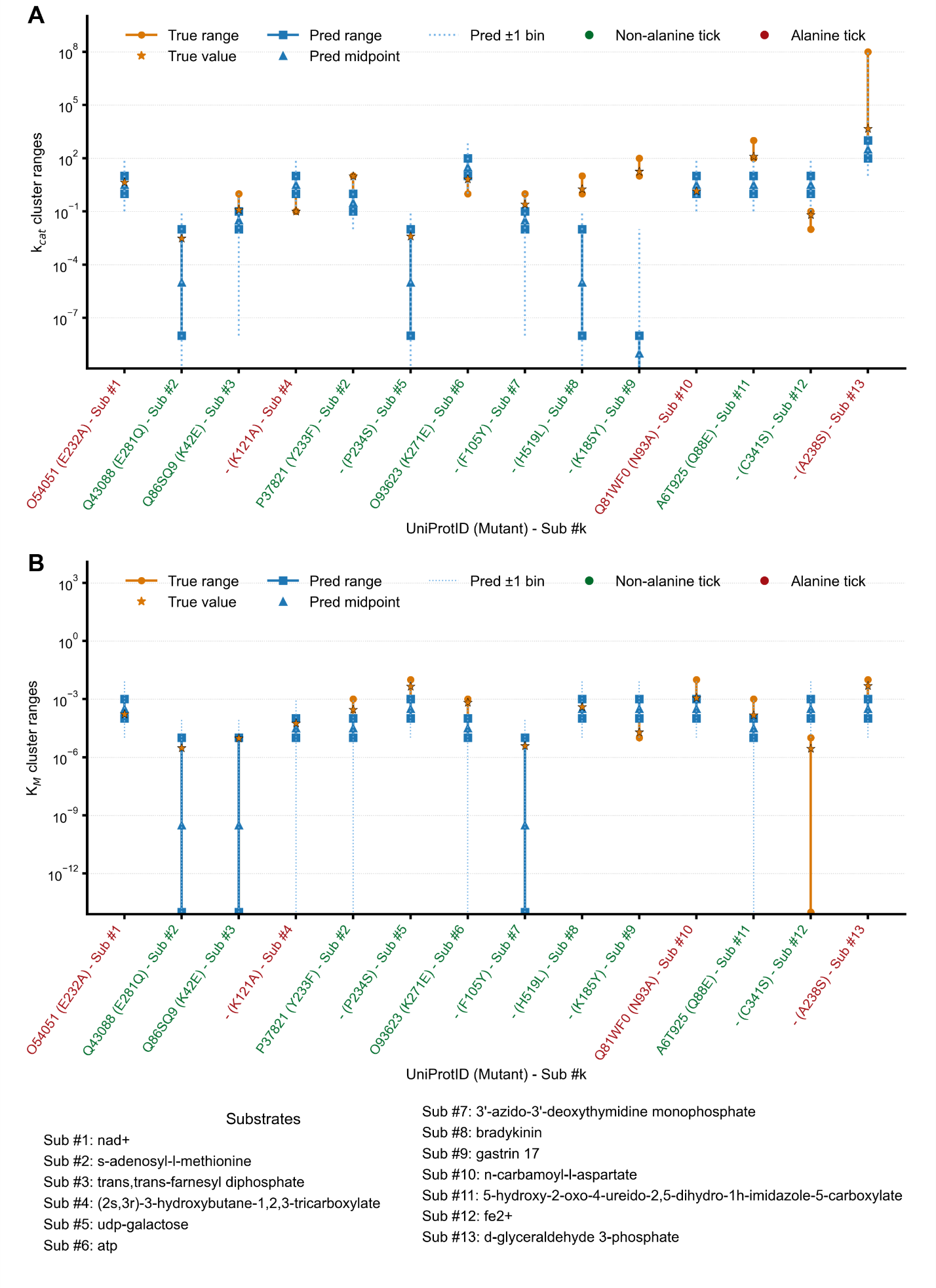
**Fig. S4. Evaluation of broader OOD mutants including non-alanine variants.**
(A) Predicted versus experimental $k_{\mathrm{cat}}$ranges for 14 enzyme–substrate pairs withheld under an ≤ 80 % sequence-identity cutoff relative to the training set. Mutants containing non-alanine substitutions (green) and alanine substitutions (red) are shown, with dashed lines representing experimental bin midpoints and solid lines denoting RealKcat predictions. (B) Corresponding analysis for $K_{M}$across the same OOD mutants, illustrating that RealKcat generalizes to substitutions beyond alanine and maintains bin-level concordance in affinity trends. Substrate identities are listed below for reference. Model performance reflects learned catalytic reasoning rather than recognition of alanine-substitution signatures, extending the generalization evidenced in Fig. 4 and 5.

#### Parameter Sharing in RealKcat

In RealKcat, a single set of hyperparameters optimized for 𝑘_𝑐𝑎𝑡_ prediction was reused for *K_M_* prediction, rather than independently re-tuning each task. This parameter-sharing strategy reflects the biochemical relationship between turnover and substrate affinity: both are shaped by the structural and functional properties of enzyme active sites and their interactions with substrates. By applying the same tuned configuration across tasks, the model leverages common patterns in the data and captures coordinated features that influence both catalysis and binding. This approach improves efficiency and reduces the risk of overfitting in relatively imbalanced datasets. However, it also introduces a trade-off: because the hyperparameters were tuned on 𝑘_𝑐𝑎𝑡_*, model performance may be more strongly aligned with turnover prediction than with affinity prediction, potentially reducing resolution for *K_M_**. Despite this limitation, the shared framework provides a principled and pragmatic means of integrating related kinetic parameters, reinforcing the interdependence of 𝑘_𝑐𝑎𝑡_ and *K_M_** while maintaining robustness and generalizability.

#### ***Catalytic Awareness of the Alkaline Phosphatase (PafA) Enzyme from Markin et al.*** (1)

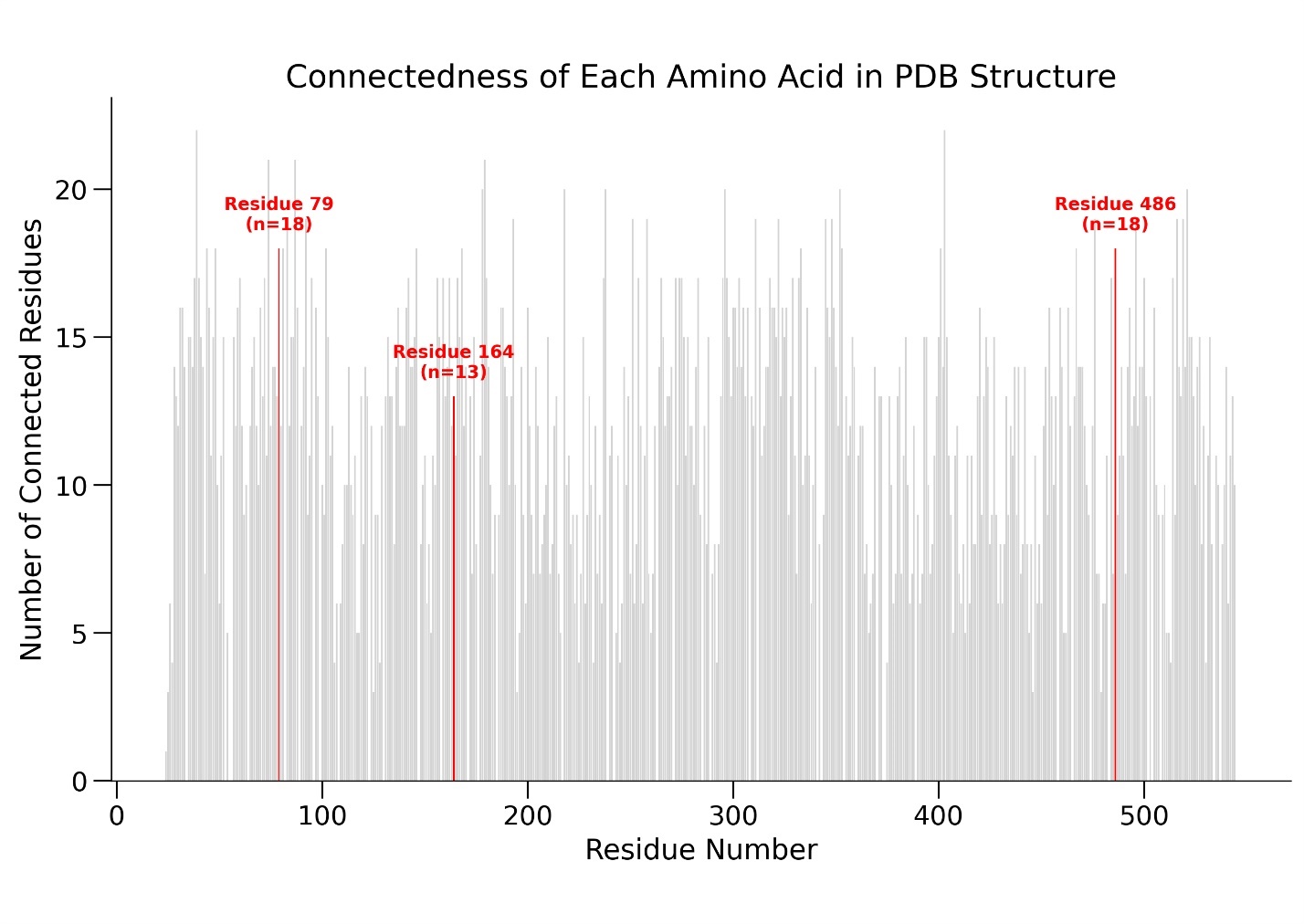

**Fig. S5. Connectivity of catalytic residues in PafA (PDB ID: 5TJ3).** This figure displays the connectivity of each amino acid residue in the PafA structure (PDB ID: 5TJ3), measured by the number of neighboring residues within a specified distance threshold of 6 Å. The gray bars represent the connectivity of each residue, with catalytic residues 79 and 164 highlighted in red. Residue 79, connected to 18 other residues, and residue 164, connected to 13, demonstrate high connectivity.

The R164 residue in PafA (PDB ID: 5TJ3) lies within a structurally significant region, as revealed by connectivity analysis (Fig. S5). With 13 neighboring contacts, R164 contributes to local structural integrity and likely stabilizes the active site. This high degree of interconnectivity suggests that substitutions at R164, particularly those altering side-chain chemistry, may disrupt local packing and perturb catalytic stability. Similar effects are observed for other highly connected residues such as H486, where experimental measurements show activity decreases exceeding two kinetic bins. RealKcat’s predictions for H486 variants (Fig. 6) highlight this sensitivity. For example, the H486V substitution, which replaces histidine with the smaller side chain of valine, was predicted by RealKcat to retain 𝑘_𝑐𝑎𝑡_ value within an order of magnitude of the WT experimental *in vivo* value. This prediction does not capture the experimentally observed destabilization, underscoring a limitation of the model in accounting for certain side-chain substitutions at structurally dense sites. The resulting misclassification points to opportunities for refinement, particularly in training RealKcat on additional valine substitutions at catalytic residues to improve its ability to model such nuanced structural effects. With the current few-shot setting, R164V is not in the training set but validation set. Overall, the structural context provided by connectivity analysis enriches interpretation of RealKcat’s predictions for PafA, illustrating both the model’s capacity to recognize mutation-sensitive regions and the areas where predictive accuracy can be further improved.
